# SYNBRIDGE: A VERSATILE FRAMEWORK FOR BIDIRECTIONAL INTERCONVERSION BETWEEN BIOBRICKS AND GOLDEN GATE ASSEMBLY STANDARDS

**DOI:** 10.64898/2026.10.16.759639

**Authors:** Alonso Molina-Calvo, Angel Navarro-Fernandez, Paola Morales, Manuel Bernal, Carlos Rodriguez-Caso

## Abstract

Component reusability and interoperability are fundamental to synthetic biology. While BioBricks laid the groundwork for part domestication, Golden Gate-based assembly standards, in particular Modular Cloning (MoClo), revolutionized multipart assembly. Despite widespread adoption, standardized conversion between these two popular frameworks remains challenging due to sequence incompatibilities. While *de novo* synthesis remains constrained by cost and sequence complexity, conventional PCR-based conversion suffers from off-target binding driven by sequence redundancy in multigene devices, unscalable primer optimization costs, and amplicon size constraints. To address interconversion from an engineering perspective, we present SynBridge: a closed-loop idempotent workflow for bidirectional BioBrick-MoClo conversion built around two core operations, Bri2Mo and Mo2Bri. As a proof of concept, we validated SynBridge by constructing a composite genetic system integrating a BioBrick module for LuxR-mediated RFP expression with a MoClo-assembled device for thermoregulated GFP expression. Quantitative cloning efficiency and functional characterization confirmed a robust and reproducible workflow that operates without requiring ad-hoc solutions. SynBridge provides a versatile engineering tool that ensures backward compatibility, maximizes part reusability, and establishes a standardized interchangeability framework suitable for automated procedures.

**GRAPHICAL ABSTRACT:** 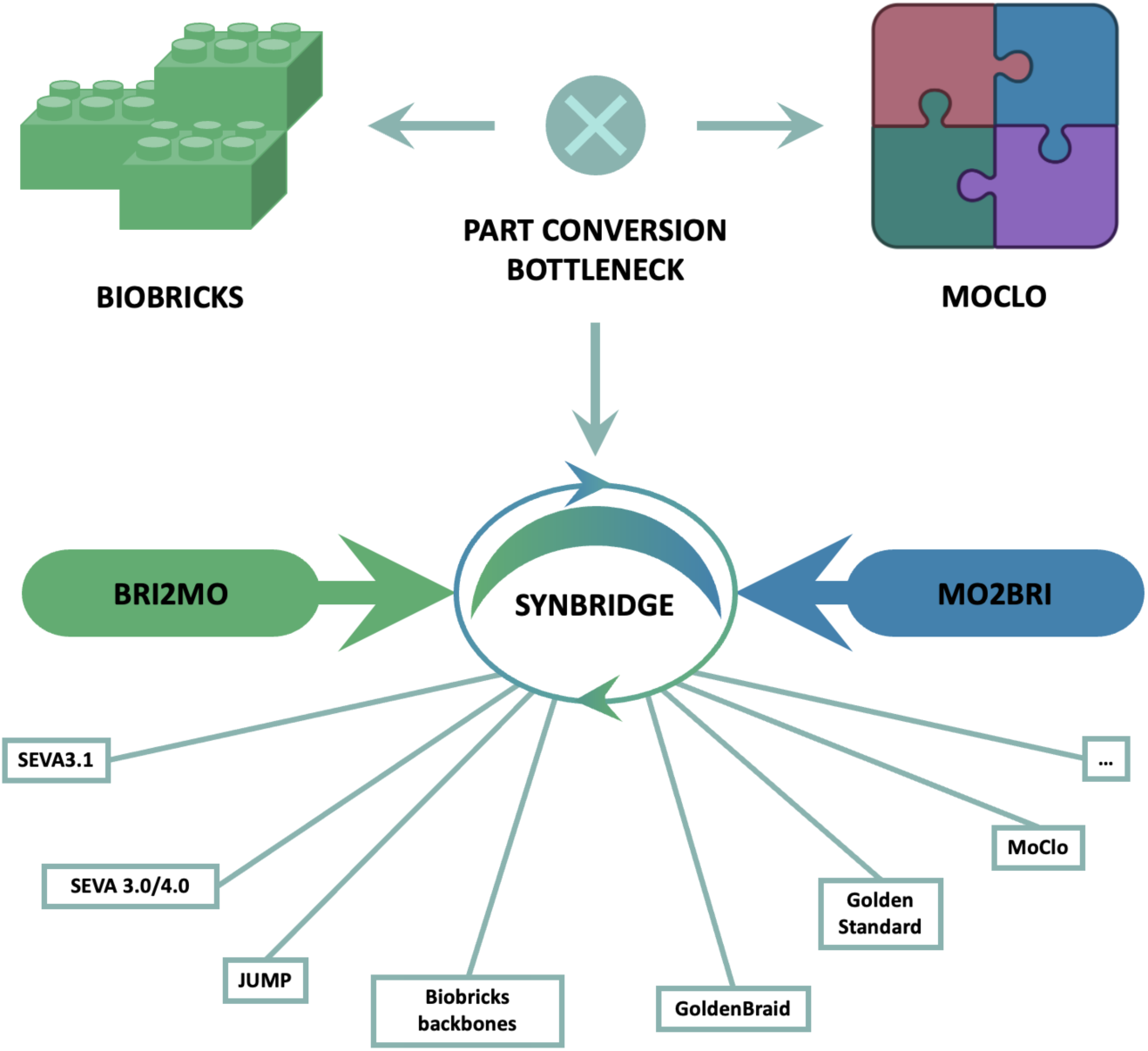

## INTRODUCTION

Synthetic Biology has driven active fundamental and applied research across a broad range of fields, from biomedicine (1) to climate change mitigation (2). Undoubtedly, major advances in DNA manipulation and genome editing have been pivotal to the development of this discipline (3, 4). However, technical innovation alone is not sufficient in itself. What distinguishes synthetic biology from any other approach is its vision of biology as a true engineering discipline—that is, an applied field grounded in a rigorous process of standardization, modular design, and rational thinking (5, 6).

A distinguishing feature of this engineering field is that components and behaviors are primarily encoded in DNA sequences (Endy, 2005). Therefore, a central focus of this discipline is the ability to write the information of genetic circuits into DNA strands (7). Despite existing limitations, advances in DNA synthesis have made it possible to obtain specific sequences on demand (8). The application of standards in design and assembly has maximized part usability (9) as it provides the community with component interchangeability. To facilitate this, the establishment of standards for effective, reliable and reusable construction of genetic parts and devices has been a central concern for a dedicated research community (10–13). Consequently, sequences—especially those that are difficult to synthesize and are meant to be conserved—can be neatly organized in repositories and DNA collections in synthetic biology labs and other repositories. This is a key aspect in formalizing the field as an engineering discipline and promoting reusability.

Traditional cloning methods, which relied on selecting appropriate enzymes to “cut and join” DNA fragments, have evolved into methodologies capable of assembling tens of fragments in a single step (Casini *et al.*, 2015). However, the major conceptual shift from traditional cloning was the emergence of idempotent methods for DNA assembly: a procedural framework where with a few rules of design and a handful of enzymes any arbitrary DNA assembly is attainable (14).

BioBrick assembly is considered one of the first idempotent DNA cloning systems (14). Its practical use requires the ‘domestication’ of genetic parts, ensuring the removal of specific DNA motifs from the sequence. The result is a framework in which a collection of compatible components is created. This standard is arguably the most widely used DNA assembly method to date, benefiting from extensive cumulative experience. Community efforts have led to the creation of the largest collection of standardized DNA parts. According to the most recent statistics, ‘The Registry of Standard Biological Parts’ (https://registry.igem.org/) contains more than 84,000 documented parts, underscoring its impact on the field.

However, this method has its own caveats. DNA parts must be assembled pairwise resulting in a labor-intensive and time-consuming process. As an alternative, the use of type IIS endonucleases by the Golden Gate assembly made possible the simultaneous assembly of multiple DNA fragments in a one-pot reaction (3). This advance gave rise to the definition of the so-called modular cloning (MoClo) (15, 16). MoClo is a Golden Gate assembly in which flanking specific genetic parts of a transcriptional unit, such as promoters, ribosome binding sites (RBS), coding sequences (CDS), and terminators are specifically designed according to a standard. These fusion sites define their position within the unit, allowing the assembly in a single reaction and in a concrete order.

Variations of the MoClo standard have been developed for particular purposes, including Golden Braid (17), EcoFlex (18), Golden Standard (19), Loop Assembly (20), Mobius assembly (21) and Phytobricks (22, 23), among others. These standards have expanded the repertoire of genetic devices and provided the community with frameworks to determine what works and what does not. From an engineering perspective, reusing tested components is essential for accelerating the Design-Build-Test-Learn (DBTL) cycle (24), particularly for transcriptional units with verified functionality or existing sequences that are difficult to obtain due to high synthesis complexity.

As widely adopted frameworks, the BioBrick and MoClo assembly standards represent key pillars in the development of synthetic biology. However, their combination in the same project is not straightforward. Efforts to enhance interoperability between standards have approached the problem from different angles. For example, the adaptation of the canonical SEVA (Standard European Vector Architecture) format to work with BioBrick was proposed as SEVA 3.1 (25), resulting in SEVAbrick. This system adapts canonical SEVA format to accept BioBrick formats in its cargo. Over this variation Damalas et al., 2020 (25) offered a methodology to interoperate between BioBrick and MoClo assemblies based on the use of a predefined primer collection and PCR amplification to convert one genetic part from one standard to another. Alternatively, the JUMP initiative (26) enables the use of both assembly standards in the same plasmid. JUMP is especially pertinent when integrating multiple transcriptional units or genetic devices that come from assemblies using varying standards into a unified framework. However, it does not facilitate their interconversion.

From a traditional perspective, joining DNA fragments with different assembly standards can be done by a custom intervention for interconversion, either via *de novo* synthesis or by a PCR-based modification of flanking sites. Suitable for isolated applications, these tailored strategies lack scalability. A common workaround for converting constructs between assembly standards involves designing custom overhang primers. In this approach, two primers per transcriptional unit (TU) anneal to the flanks of the sequence—bypassing the original standard’s prefix and suffix—while their 5′ overhangs introduce the features of the new assembly method. However, this strategy presents critical limitations that preclude it from serving as a universal solution desirable for standardized or highly automated workflows. First, multigene devices often rely on a limited set of terminators, and promoters are frequently reused across TUs; such sequence redundancy leads to off-target primer binding, invalidating specific PCR amplification and downstream site-directed mutagenesis. Second, scaling this process across a repository creates a substantial burden in terms of time and cost, as every fragment requires custom primers and PCR optimization. Finally, synthetic biology is scaling up from multi-TU constructs to minimal synthetic genomes (27). In this context, PCR-based conversion is constrained by amplicon size limits, as amplifying constructs larger than ∼5 kb typically fails without complex, specialized protocols.

From an engineering perspective a desirable design for format interconversion must preserve device integrity, allowing for an ideally unlimited number of conversions with no alteration in the sequence and no variation in the assembly rules. Our proposed framework achieves this by enabling an idempotent interconversion between the two standards. This approach is closely related to the concept of ‘round-tripping’, a systems engineering term (28), which refers to the ability to convert an artifact from one format to another and back again without losing integrity or degrading the original data.

To address these limitations, we present a rational design strategy that establishes a lossless, closed-loop workflow for idempotent conversion between BioBrick and MoClo assemblies. This framework allows transcriptional units to be indefinitely interchanged without compromising sequence integrity, while enabling the incorporation of new genetic devices to expand functionality, interoperability, and reusability.

## MATERIAL AND METHODS

### Molecular Biology reagents

All synthetic DNA sequences were provided by Integrated DNA Technologies (IDT, Coralville, IA,USA). General molecular biology reagents, including CaCl2 molecular-grade water and Luria-Bertani (LB) broth, were obtained from Sigma-Aldrich (St. Louis, MO, United States). Restriction enzymes, recombinant shrimp alkaline phosphatase (rSAP), T4 DNA ligase, *EcoR*I-HF, *PstI*-HF, *Xba*I, *Spe*I-HF, *Not*I-HF, *BsmB*I-v2, *Bsa*I-v2, *Eco*RV and NEB 10-beta competent *Escherichia coli* cells were supplied by New England Biolabs (NEB, Ipswich, MA, USA). Taq DNA polymerase was purchased from PCR Biosystems (London, UK).

Golden Gate assemblies were performed as previously described by (37). Plasmid DNA was extracted using the Quick-DNA™ Miniprep Plus Kit (Zymo Research, Irvine, CA, USA) according to the manufacturer’s instructions, and DNA concentrations were determined using a NanoDrop One spectrophotometer (Thermo Fisher Scientific, Waltham, MA, USA). PCR products and DNA fragments were purified using the NucleoSpin Gel and PCR Clean-up kit (Macherey-Nagel, Düren, Germany). Bacterial transformation cells were carried out using CaCl2 chemically induced competents from *E. coli* NEB 10-beta (see Supplementary Data for methodological details). Sequence verification of the assembled constructs was outsourced to STAB VIDA (Caparica, Portugal) and GENEWIZ (South Plainfield, NJ, USA).

### Assembly of plasmid collections

To construct the MCS1-4 into pSB1A3, pSB1C3, and pSB1K3 plasmid collections, MCS with the DO (Bba_J44050) sequence was ordered to IDT (Figure 1B) with EcoRI and PstI flanking sequences with four base pair extra outwards the sequence for optimal endonuclease activity. Acceptor vectors were obtained from the part registry. MCS insertion of MCS into acceptor backbones was done by digestion with *EcoR*I and *Pst*I (Figure 1A). Subsequent ligation reactions were performed utilizing a 3:1 insert-to-backbone molar ratio. Ligation products were transformed into competent NEB 10-beta cells, and successful transformants (see protocol in Supplementary Data) were isolated and verified via colony PCR followed by whole-plasmid sequencing (GENEWIZ).

**Figure 1.**
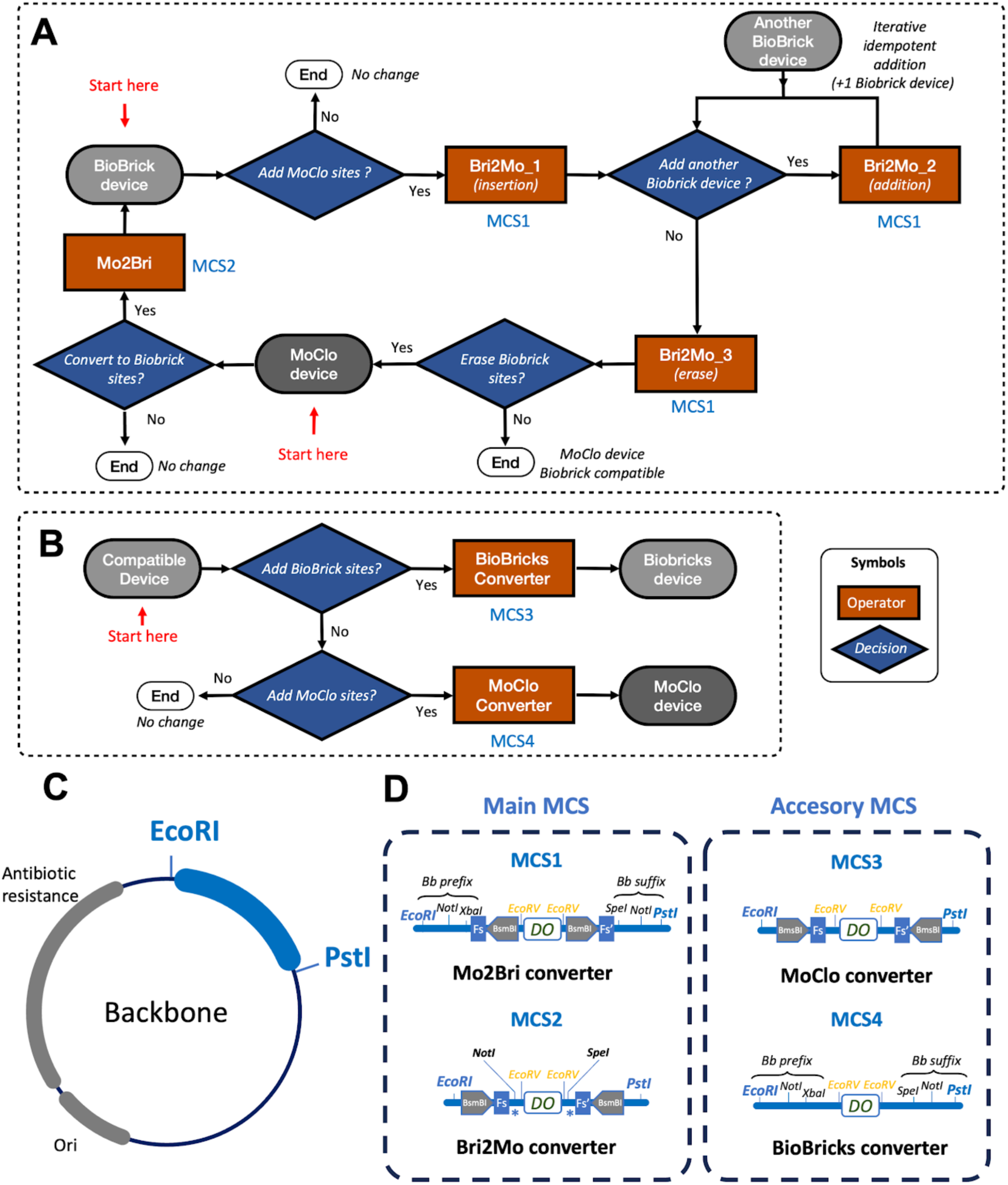
SynBridge workflow for closed-loop, idempotent, bidirectional conversion between BioBricks and MoClo assembly standards. (A) Orange rectangles represent the operators detailed in Figures 3 and 4, while blue diamonds represent the decision nodes. MCS1 and MCS2 indicate the specific multiple cloning sites (MCS) associated with each operator (see Figure 2 for details). Accessory workflow (B). For the sake of completeness, an additional process with its respective MCS is included in the system to facilitate the incorporation of non-standardized devices into BioBricks or MoClo standards; here, MCS3 and MCS4 indicate the multiple cloning sites involved with the operators for the respective BioBricks and MoClo conversions (see Figure 2 for details). A “compatible device” is defined as any DNA part that does not contain within its sequence the BioBrick prefix and suffix restriction sites (EcoRI, SpeI, XbaI, PstI, and NotI), nor the MoClo Type IIs recognition sites (specifically BsaI, as well as other sites used in higher assembly levels such as BsmBI, as proposed in this work). Ensuring the absence of restriction sites for enzymes involved in both BioBricks and MoClo guarantees the full compatibility of the newly added devices with SynBridge. Architectural design and constraints of the SynBridge Multiple Cloning Sites (MCS). (C) Structural requirements for a SynBridge-compatible backbone. The MCS must be flanked by an upstream EcoRI and a downstream PstI restriction site located within a non-essential vector region. Additionally, the backbone must be completely devoid of *Spe*I, *Not*I, *Bsa*I, *BsmB*I, and *EcoR*V recognition sequences. (D) Architectures of the primary and accessory MCSs. MCS1 (Mo2Bri) and MCS2 (Bri2Mo) mediate the bidirectional interconversion between the BioBricks and MoClo standards through Mo2Bri and Bri2Mo operations. As an additional functionality, the insertion of two flanking cutting outwards BsaI recognition sites allow for using MCS2 as an acceptor for level 1 MoClo assemblies. Accessory cassettes MCS3 and MCS4 facilitate the direct integration of synthesized or PCR-amplified DNA fragments via EcoRV blunt-end cloning. All designs incorporate a phenotypic dropout (DO) reporter for negative selection. Abbreviations: *, BsaI restriction site; Bb, BioBricks; Bri2Mo, BioBricks to MoClo; DO, dropout reporter; Fs, GGAG; Fs’, CGCT; Mo, MoClo; Mo2Bri, MoClo to BioBricks; ori, origin of replication. See supplementary Table S1 to get access to MCS sequences.

### Mo2Bri operation

Mo2Bri operation was performed using MCS1 (Figure 1B) applying one of three distinct cloning strategies: (i) One-pot restriction-ligation Golden Gate assembly. The protocol used was obtained by (37). It was employed, modifying the thermal cycling parameters to include digestion steps either at 37 °C or 42 °C to optimize the efficiency of the Type IIS restriction enzyme *BsmB*I (see protocol in Supplementary Data). (ii) Sequential restriction and ligation. A two-step reaction was performed, consisting of an initial *BsmB*I digestion followed by a separate ligation step utilizing a 3:1 insert-to-vector molar ratio. (iii) Sequential restriction-ligation with vector dephosphorylation. This approach followed the two-step protocol described above, with the inclusion of an intermediate step prior to ligation where the digested acceptor backbone was treated with rSAP to prevent self-circularization. For further technical information see Supplementary Data.

### Bri2Mo operation

Bri2Mo operation was performed using MCS2 (Figure 1B). It was conducted through two sequential cloning steps. According to Figure 3A step Bri2Mo_1 consisted of digesting both the first BioBricks unit and the Bri2Mo acceptor vector with *Not*I and *Spe*I. The acceptor backbone was subsequently dephosphorylated using rSAP prior to ligation, maintaining a 3:1 insert-to-vector molar ratio. In step Bri2Mo_2, the intermediate plasmid generated in Bri2Mo_1 served as the new acceptor, which was digested with *Not*I and *Xba*I. Following dephosphorylation with rSAP, it was ligated with a second *Not*I/*Spe*I-digested BioBricks device under the same 3:1 molar ratio condition. For further technical information see Supplementary Data.

Following assembly, residual restriction recognition sites were eliminated from the backbones via site-directed mutagenesis by Bri2Mo_3. This backbone purging was achieved through PCR amplification using a series of universal primers designed to introduce specific point mutations, thereby ablating the remaining *Not*I, *Xba*I, and *Spe*I sites in the Bri2Mo_3 constructs. For further technical information see Supplementary Data.

### Image acquisition and automated phenotypic analysis

The integration of smartphone imaging with machine learning-based computer vision has recently emerged as a highly robust and reproducible standard for automated high-throughput colony enumeration and classification (38, 39). Prior to imaging, *E. coli* cultures were serially diluted and plated to achieve a statistically valid range of 30 to 300 colonies per plate. The resulting agar plates were subsequently imaged using an iPhone 16 Pro Max smartphone camera while illuminated on a UV transilluminator to excite fluorescent reporters. Image processing, colony segmentation, and phenotypic quantification were performed using SPECTRA (Spectral Phenotyping Engine for Colony Tracking and Rapid Analysis), a custom Python-based computer vision pipeline developed in this study. Detailed information about its construction can be found in Supplementary Data. This code is available for the community at GitHub (https://github.com/alonso1310/SPECTRA).

### Qualitative and quantitative fluorescence analysis

For robust phenotypic validation, 90 independent colonies per construct were picked and cultured in pairs of LB-agar plates containing the appropriate antibiotic. For each pair, one of the plates contained 3-oxohexanoyl homoserine lactone (3OC6HSL) (MilliporeSigma, Burlington, MA, USA.) at a concentration of 1 µM, which allowed the induction of the luxR mediated RFP expression device. Afterwards, each of these colonies was inoculated into a 96-well microplate containing LB broth supplemented with the appropriate antibiotics and 1 µM of 3OC6HSL. The microplates were incubated in an Agilent BioTek Synergy H1 multimode reader (Santa Clara, CA, USA) at 37 °C with continuous orbital shaking (282 cpm, 3 mm amplitude) for a 24-h kinetic runtime. Measurements were recorded at 10 minute intervals. To monitor cell growth, optical density was measured via absorbance at 660 nm (OD660) according to protocol described in (40). Simultaneously, fluorescence expression was quantified using bottom optics (read height: 7 mm) and a high-energy Xenon flash lamp. Green fluorescent protein (GFP) was detected at an excitation/emission of 479/520 nm, while red fluorescent protein (RFP) was measured at 586/611 nm; the detector gain was strictly set to 75 for both fluorescent reporters. To calculate the relative fluorescence units (RFU), raw fluorescence and OD660 values were first background-subtracted using sterile LB media blanks. The corrected fluorescence intensities were then normalized against their corresponding corrected optical densities, explicitly extracting the final reported values from the kinetic time point corresponding to the maximum expression peak of each construct. For graphical clarity, the presented figures illustrate the RFU of three representative biological replicates, whereas the comprehensive raw dataset encompassing all 90 evaluated colonies is provided in the Supplementary Data.

## RESULTS

### Closed-loop Workflow Design

Figure 1A illustrates the closed-loop workflow, which comprises two core functionalities. The first enables transcriptional units to undergo round-trip conversion—from BioBrick to MoClo and vice versa—while preserving their sequence integrity and functionality, regardless of the target standard. The second one allows additional BioBrick transcriptional units to be introduced during BioBrick-to-MoClo conversion in an idempotent manner.

Bri2Mo, as illustrated in Figure 1A, performs this BioBrick-to-MoClo conversion and consists of three distinct operations. The first operator, Bri2Mo_1, is responsible for introducing a BioBrick device into a MoClo system giving rise to a MoClo-BioBrick hybrid. The second operator, Bri2Mo_2, enables an idempotent integration of additional BioBrick devices following similar BioBrick assembly logic as it shall be detailed later. The third operator, Bri2Mo_3, eliminates the BioBrick restriction sites to yield a standard MoClo-compatible device. Additionally, Figure 1B introduces two accessory processes that permit the incorporation of novel DNA parts with no flanking adaptors into either standard. To ensure full compatibility, these sequences must lack any restriction sites for the enzymes involved in both the BioBrick and MoClo standards. Given the diversity of existing BioBrick and MoClo variants, a universal, one-size-fits-all approach is not a trivial task. In this work, we applied this exclusion criterion as the RFC-10 BioBrick standard does, which is the most abundant variant in the Registry of Standard Biological Parts. Regarding MoClo, to maintain compatibility with hierarchical, idempotent systems, we exclude *Bsa*I recognition sites, as this enzyme is utilized at Level 1 assembly in the most MoClo frameworks (see Table 1 of a compilation of standards based on the golden-gate assembly). However, the greater heterogeneity of the second Type IIs enzyme across different platforms makes it difficult to have an unique solution for all of them. In our case, we selected to avoid *BsmBI* sites. In this way, we allow compatibility with at least some of the most prominent hierarchical, idempotent MoClo assemblies, platforms and backbone standards. This is the case of GoldenBraid 2.0 (29), MoClo (15) and EcoFlex as well as with the JUMP system (26). See Table 1 for compatibility details with standards).

**Table 1.** Compatibility of the SynBridge system with existing DNA assembly frameworks and vector standards.

| Standards | Key enzymes involved | Donor devices from the standard** | Synbridge operator compatibility | Reference |
| --- | --- | --- | --- | --- |
| Loop Assembly | *Bsal*-*SapI* | L1 TU (*SapI*) | Mo2Bri (*BsmBI*) | (20) |
| Golden Braid, EcoFlex | *Bsal*-*BsmBI* | L1 TU (*BsmBI*) | Mo2Bri (*BsmBI*) | (18, 29) |
| Golden standard (SEVA), MoClo, CIDAR, Phytobricks | *Bsal*-*BpiI* | L1 TU (*BpiI*) | Mo2Bri (*BsmBI*) | (15, 19, 22, 23, 30) |
| Mobius | *Bsal*-*AarI* | L1 TU (*AarI*) | Mo2Bri (*BsmBI*) | (21) |
| JUMP* | *EcoRI*, X, S, N, *PstI*, *BsmBI* | L1 TU (*BsmBI*)<br>Bb_device (N+S)<br>Bb_device (N+S) | Mo2Bri (*BsmBI*)<br>Bri2Mo_1 (N+S)<br>Bri2Mo_2 (N+X) | (26) |
| SEVA 3.1* | N, *EcoRI*, X, S, *PstI* | Bb_device (N+S)<br>Bb_device (N+S) | Bri2Mo_1 (N+S)<br>Bri2Mo_2 (N+X) | (25) |
| pSB1C3*, pSB1K3, pSB1A3*, pSB1C5* | N, *EcoRI*, X, S, *PstI* | Bb_device (N+S)<br>Bb_device (N+S) | Bri2Mo_1 (N+S)<br>Bri2Mo_2 (N+X) | The Registry of Standard Biological Parts |

As an additional requisite to maximize interoperativity among a larger number of available backbones from the community we define a structural requirements for any multicloning site (MCS) involved in this work. As a general rule any MCS must be flanked by a *EcoR*I and *Pst*I (see Figure 1C) as it happens in Biobricks standard, SEVA 3.1 and JUMP do with their respective cargos. According to the abovementioned restrictions, Table 1 shows SynBridge operation compatibility with popular backbone systems used in synthetic biology. Figure 1D indicates the scheme of the MCS defined in this work. To expand cloning options, the dropout (DO) cassette is flanked by *EcoR*V sites, facilitating direct, blunt-end cloning of DNA fragments prepared via PCR amplification or chemical synthesis. As we shall detail later, an additional functionality that completes the interoperativity of this platform, MCS2 works as an acceptor for level 1 *Bsa*I-MoClo assemblies. This is due to it containing two flanking *Bsa*I recognition sites cutting outwards as detailed in Figure 1D. MCS sequences are indicated in Table S3 in Supplementary Data.

### Mo2Bri operation: the MSC1

Figure 2A details the workflow by Mo2Bri operation. MCS1 permits the conversion of a Level 1 MoClo assembled into a BioBricks format. Our method takes advantage of the common syntax overhangs adopted for basic part fusion sites at Level 0 (22). Consequently, any transcription unit (TU)— regardless of the Golden Gate standard used—is flanked by identical fusion sites and inward-oriented Type IIS recognition sites. Specifically, these consist of the 5’–GGAG–3’ site (Fs, upstream of the TU) and the 5’–CGCT–3’ site (Fs’, downstream of the TU). As a result, assembled TUs generated by major frameworks share these universal Fs and Fs’ boundaries. This is the case of MoClo (15), GoldenBraid 2.0 (29), Golden Standard (19), Phytobricks (23), EcoFlex (18),Yeast toolkit (13), CIDAR (30) among others.

**Figure 2.**
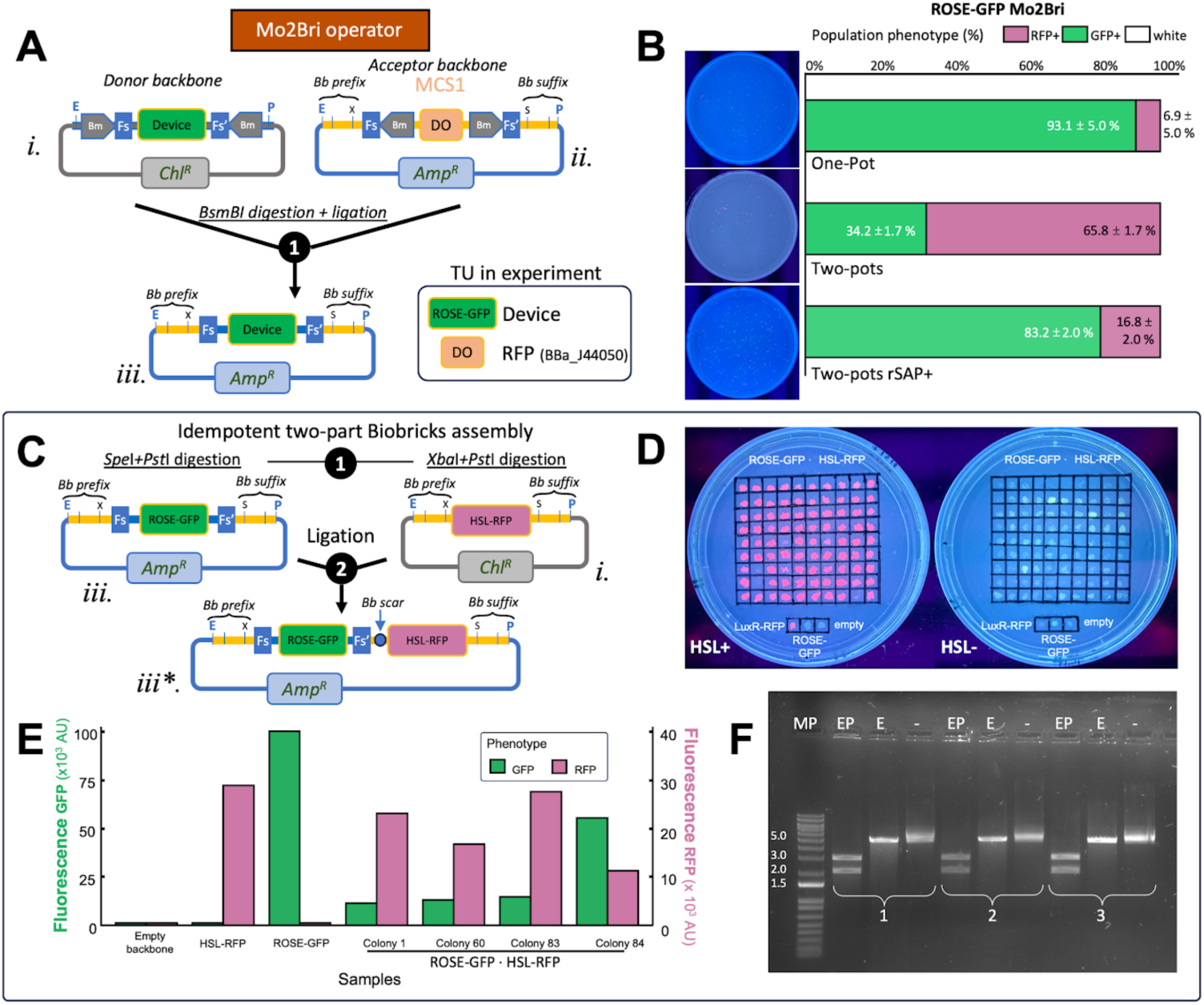
Mo2Bri operation: conversion from BioBricks to MoClo. (A) Schematic representation of the Mo2Bri operation. A MoClo device provided by a backbone Level1 MoCLo acceptor backbone. Accordingly, the device of interest must be flanked by the fusion sites of any TU sites (that we call Fs and Fs’) followed by a BsmBI site inwards. Device must be integrated into an acceptor plasmid that contains MCS1 via a BsmBI-ligation reaction, displacing a dropout (DO) reporter. The box inlet indicates the names of the devices involved in the proof of principle of this procedure: ROSE-GFP, a temperature-dependent inducible GFP expression system was introduced in a MSC1 with a BBa_J44050, an RFP gene controlled by Plac wring as the DO. **(B)** Percentage of the Mo2Bri operation of ROSE-GFP initially obtained by its respective basic parts assembling according to MoClo (see supplementary data for the description of this TU). Phenotypic screening by colony counting shows cloning efficiencies across protocols, with one-pot conditions yielding the highest efficiency. Percentages are indicated as the median ± standard deviation from three independent biological replicates starting from independent cloning reactions. **(C)** Schematic representation of the two-part BioBrick assembly demonstrating that the resulting converted construct operates as a BioBrick part. The scheme indicates that ROSE-GFP after Mo2Bro operation (green colony phenotype under the tested temperature condition) was fused to LuxR-RFP. **(D)** Phenotypic validation of inducible behavior was performed by replica plating the 90 colonies resulting from the cloning described in the previous panel. The plate on the left lacks HSL in the medium, while the plate on the right contains 1 µM HSL. **(E)** Spectrofluorometric plate-reader measurements were used to quantify red and green fluorescence normalized to optical density at 660 nm for each of the 90 colonies over a 24 h period (single experiment per colony). The graph shows the representative behavior of fused devices compared to the constructs alone. **(F)** DNA gel electrophoresis of the restriction digest confirmed the expected band sizes (bottom right). Three randomly chosen colonies were cultured for plasmid miniprep isolation and subsequent enzymatic digestion (MP: Molecular pattern DNA ladder; EP: EcoRI/PstI double digest; E: EcoRI single digest). E/P digestion yielded a band of 2930 bp corresponding to the sum of ROSE-GFP (1014 bp) + LuxR-RFP (1916 bp), alongside the empty acceptor plasmid pSB1A3 used in the experiment (2029 bp). The digestion with *EcoR*I exhibited a single band corresponding to the expected sum of the plasmid backbone and construct sizes.

In MoClo (and also in Golden Braid and EcoFlex) L1 destination backbone provides two *BsmBI* recognition sites to operate in the L2 over Fs and Fs’ sites, respectively. This is represented by plasmid *(i)* in Figure 2A. Plasmid *(ii)* contains MCS1 with biobricks sites and a *BsmB*I oriented outwards in their cargo. In the same Figure 2A, the result of a *BsmBI* digestion and further ligation (step 1) for constructions (i) and (ii) is represented by plasmid (iii).

Figures 2B and 2C illustrate the experimental proof of concept, providing quantitative measurements of process efficiency. To this end, we used a previously assembled MoClo-built termoregulatable GFP expression device controlled by a Repression Of heat Shock gene Expression or ROSE (31) (hereafter ROSE-GFP). The device was obtained from their basic parts doing a one-pot *Bsa*I MoClo reaction. ROSE domestication to be compatible with the MoClo standard is described in SI. The Level 1 acceptor backbone was pSB1C3·MSC2, since took advantage of the abovementioned additional functionality of MCS2 (Figure 1D) when it is cut with *Bsa*I. MCS2 was introduced in pSB1C3 by *EcoR*I and *Pst*I digestion. Accordingly, ROSE-GFP in pSB1C3·MSC2 corresponds with the donor construct (i) in the general scheme in Figure 2A. DNA sequencing confirmed that the MoClo ROSE-GFP device was flanked by *BsmB*I with Fs and Fs’ fusion sites. The backbone acceptor was constructed for experimental validation using a pSB1C3 with MSC1 also between the *EcoR*I and *Pst*I sites (named as pSB1C3·MCS1). In the experimental validation, MCS1 contained as a dropout (DO) the BBa_J04450, encoding a constitutive RFP production. *BsmB*I digestion of donor (i) and acceptor backbones (ii) and further cloning were done as illustrated in step 1 in Figure 2A. Three different cloning variants were done in order to evaluate an optimisation of the process.

Figure 2B shows the experimental results of the different variants. The first condition was done on a sequential *BsmB*I digestion-ligation (named as Two-Pot condition). It inserts the MoClo assembled ROSE-GFP into the pSB1C3·MSC1 resulted in an irreversible conversion to a Biobricks format. The process yielded 34.2 ± 1.7% correct transformants. Green fluorescence under UV light during incubation and under induction conditions, i.e. 37°C, confirmed that the Mo2Bri operation did not alter the expected phenotypic behaviour. The yield of positive colonies increased to 83.2 ± 2.0% with the addition of alkaline phosphatase (Two-Pot rSAP+). In order to simplify the procedure we tested a One-Pot variant (see methods section for details). The optimisation resulted in a statistically significant increase of efficiency of 93.1 ± 5.0% respecting Two-Pot SAP+ condition (t-student, p-value < 0.05). These results came from three independent biological replicates starting from independent cloning reactions. Sequencing of three clones selected from the independent replicates for the more successful One-Pot protocol confirmed 100% sequence identity with the theoretical sequence, indicated in Table S2 in Supplementary Data.

To confirm that the ROSE-GFP TU, hosted in pSB1C3-MCS1, was functional as a Biobricks device, a *Spe*I/*Pst*I, *Xba*I/*Pst*I two backbone Biobricks assembly variant was performed (see Figure 2C). The Biobrick assembly was done with a BioBricks device codifying a previously built homoserin lactone (HSL)-inducible expression of a RFP mediated by LuxR, hereafter referred to as HSL-RFP. See details of its construction in Supplementary Data. The result was a tandem of ROSE-GFP followed by HSL-RFP as confirmed by sequencing of three independent clones with 100% sequence matching the theoretical sequence. This confirmed that *BsmB*I recognition sites were absent from the sequence and conveniently replaced by BioBricks sites, ready for another idempotent Biobrick iteration. Figure 2D shows the UV-illuminated plate screening confirming the expected behavior when 37°C GFP and HSL addition were applied to the joint construct. Figure 2E showed that randomly selected colonies had expected phenotypic behaviour in liquid cultures when induced by temperature (GFP expression) and HSL (RFP expression). Figure 2F confirmed restriction mapping giving the expected digestion profiles.

### Bri2Mo operation: the MSC2

As Figure 1A shows, the Bri2Mo workflow is divided into three differentiated operation: Bri2Mo_1 for adding a new Biobrick device, Bri2Mo_2 to provide an idempotent process to add additional devices at this point of the conversion and, Bri2Mo_3 for erasing Biobricks marks. All of them are made by using MCS2 (see Figure 1B for the scheme and supplementary Table S1 for the sequence). Accordingly, Figure 3 details the cloning process and the experimental validation for the migration of a BioBricks device into a MoClo level 1 unit.

**Figure 3.**
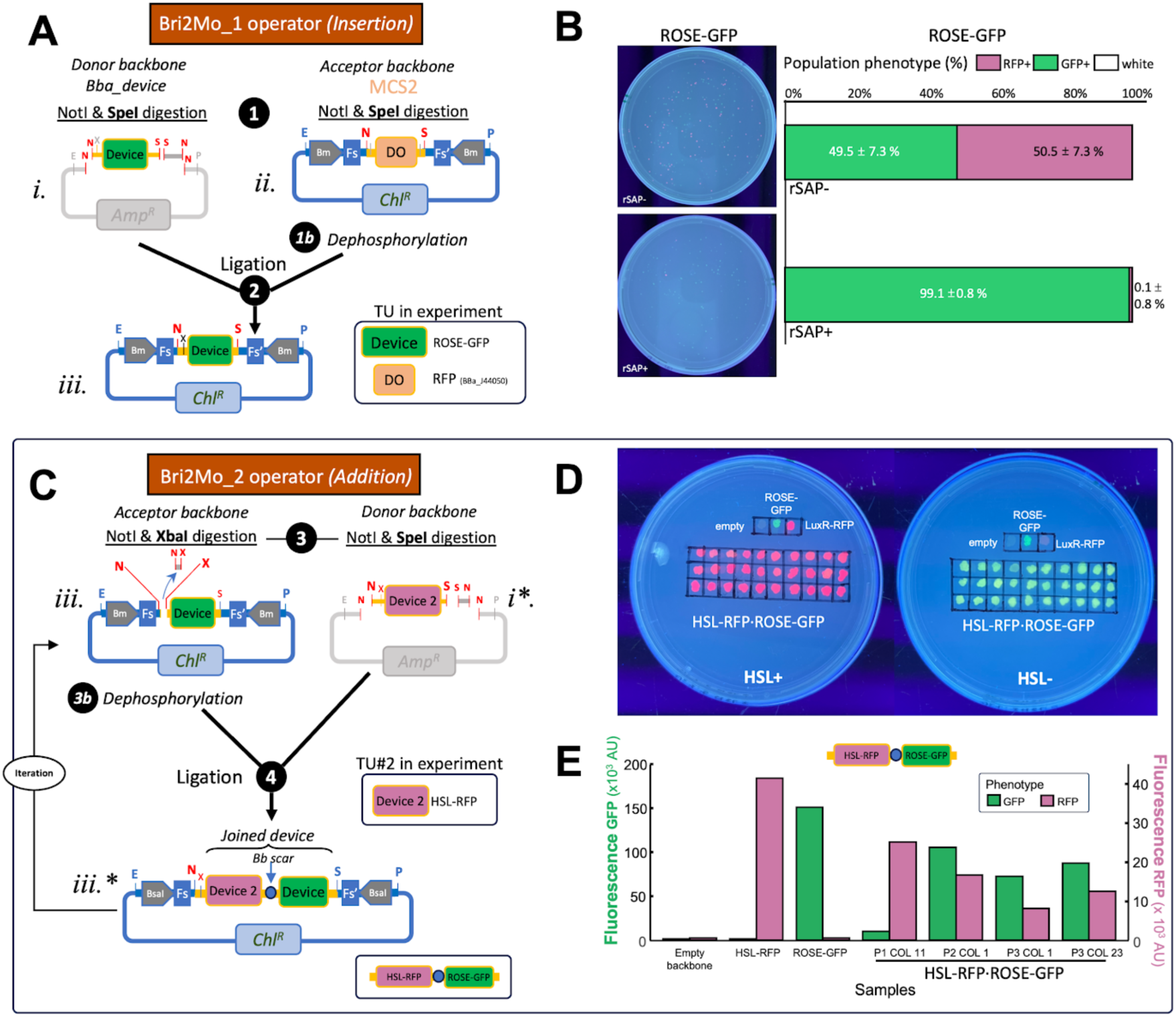
Bri2Mo operation: conversion from MoClo to BioBricks. (A) Schematic representation of the Bri2Mo_1 operation for inserting a BioBrick part into MCS2. A BioBrick device hosted in a donor backbone is transferred to MCS2 located in a compatible acceptor backbone. Step 1 represents NotI/SpeI digestion of both backbones, followed by ligation (Step 2). Dephosphorylation of the acceptor backbone is indicated in Step 1b. **(B)** Experimental validation of the assembly efficiency described in panel A. A ROSE-GFP BioBrick device was cloned into MCS2 containing BBa_J44050 acting as a dropout reporter (DO, red fluorescence). Comparing the non-dephosphorylated protocol (rSAP–) with acceptor backbone dephosphorylation (Step 1b, rSAP+) demonstrated that rSAP treatment notably increased cloning efficiency. No white colonies were observed without dephosphorylation. **(C)** Schematic representation of the Bri2Mo_2 operation (idempotent addition). The sequential addition of a second BioBrick device is achieved by ligating a NotI/XbaI-digested acceptor with a NotI/SpeI-digested insert. **(D)** Functional validation of the HSL-RFP·ROSE composite construct generated by the Bri2Mo_2 operation. All transformants resulting from cloning were replica-plated onto media with 1 µM HSL (HSL+) and without HSL (HSL–). **(E)** Spectrofluorometric quantification of randomly picked colonies of a liquid culture with 1 µM HSL (HSL+).

Bri2Mo_1 operation has the function of *insertion*: a BioBricks device (donor plasmid (i), Figure 3A) is inserted into a compatible MCS2 backbone (acceptor plasmid (ii), Figure 1A) via *Not*I and *Spe*I double digestion, heat inactivation, and ligation. It is worth stressing that MCS2 lacks a *Not*I site in the suffix. MC2 permits the insertion of a BioBrick device in a way that permits further idempotent addition of Biobricks devices by means of an non-cannonical BioBricks assembly. In the Bri2Mo_1 process, plasmid (i) releases a short *Not*I/*Spe*I fragment, while acceptor plasmid (ii) releases the DO. A dephosphorylation (step 1b) is an optional step that prevents acceptor plasmid (ii) recircularization.

In our experimental validation *Not*I and *Spe*I double digestion in Bri2Mo_1 replaced the fluorescence dropout (DO) reporter (BBa_J04450, encoding RFP) in the MCS2 acceptor vector with the device of interest: the BioBricked ROSE-GFP already converted by Mo2Bri operation described in the previous subsection (see box inlet in Figure 3A). Supplementary Data provides a detailed protocol and the DNA involved sequences.

Assembly efficiency for the initial insertion made by Bri2Mo_1 operation was evaluated by phenotypic colony screening after transformation (Figure 3B). A digest-ligation approach without acceptor dephosphorylation (rSAP-) yielded 49.5% ± 7.3% correct transformants (GFP+). The addition of alkaline phosphatase (rSAP) significantly increased assembly efficiency to 99.1% ± 0.8% (t-student, P<0,05). Attending this cloning efficiency the 11 nucleotide *Not*I/*Spe*I fragment released from the donor backbone did not appear to interfere in the cloning efficiency. No white colonies were detected. Sequencing of three independent clones from this step confirmed 100% sequence identity with the theoretical sequence, indicated in Table S2 in Supplementary Data.

As the new condition of the device plasmid (iii) provides the property of the addition of new devices in an idempotent way by means of a Bri2Mo_2 operation. This was evaluated through the incorporation of a second BioBrick device (Figure 3C). The acceptor plasmid resulting from the previous step was prepared by *Not*I/*Xba*I digestion, while the incoming device was prepared by *Not*I/*Spe*I digestion. This non-cannonical biobricks operation generates a scar-mediated assembly. The idempotency of this operation is a consequence of the modification of the Biobrick suffix. Plasmid (iii) and (iii*) in Figure 3 reveals the absence of *Not*I in the suffix and it explains the idempotency of this non-cannonical biobrick.

To experimentally verify this idempotency of the Bri2Mo_2 operation, the HSL-RFP device previously used in the previous subsection was assembled into the plasmid (iii) with ROSE-GFP obtained applying Bri2Mo_1. Figure 3D shows the phenotypic behaviour of all transformants by applying replica plating of presence or absence of HSL (controlling the RDF expression) and incubating at 37°C (activating GFP expression). Additionally, the sequencing of three independent clones from this assembly step confirmed 100% sequence identity with theoretical expected sequences. Figure 3D spectrofluorometric assays and UV-illuminated plate screening showed the constitutive expression of GFP alongside RFP induction dependent on the addition of HSL while Figure 3E shows the behaviour in liquid cultures.

Finally, Figure 4A shows the process of Bri2Mo_3, responsible for erasing all BioBricks sites and to generate a MoClo-compatible DNA device. Erasing process eliminates restriction recognition sites from the backbones via site-directed mutagenesis through PCR amplification using a series of standardized primers, named as Erasing forward and reverse primers (see Supplementary Table S2). PCR amplification uses as the template the backbone with MCS2 with the donor fragment inserted. Specific primer recognition gives a DNA amplification ablating *Not*I, *Xba*I, and *Spe*I sites. Worth to stress, they have been designed to hybridize with the sequence related to the MCS2. This avoids the need to design specific primers for the erasing of any newly assembled part, providing a general, one-for-all process for any piece following Biobricks rules.

**Figure 4.**
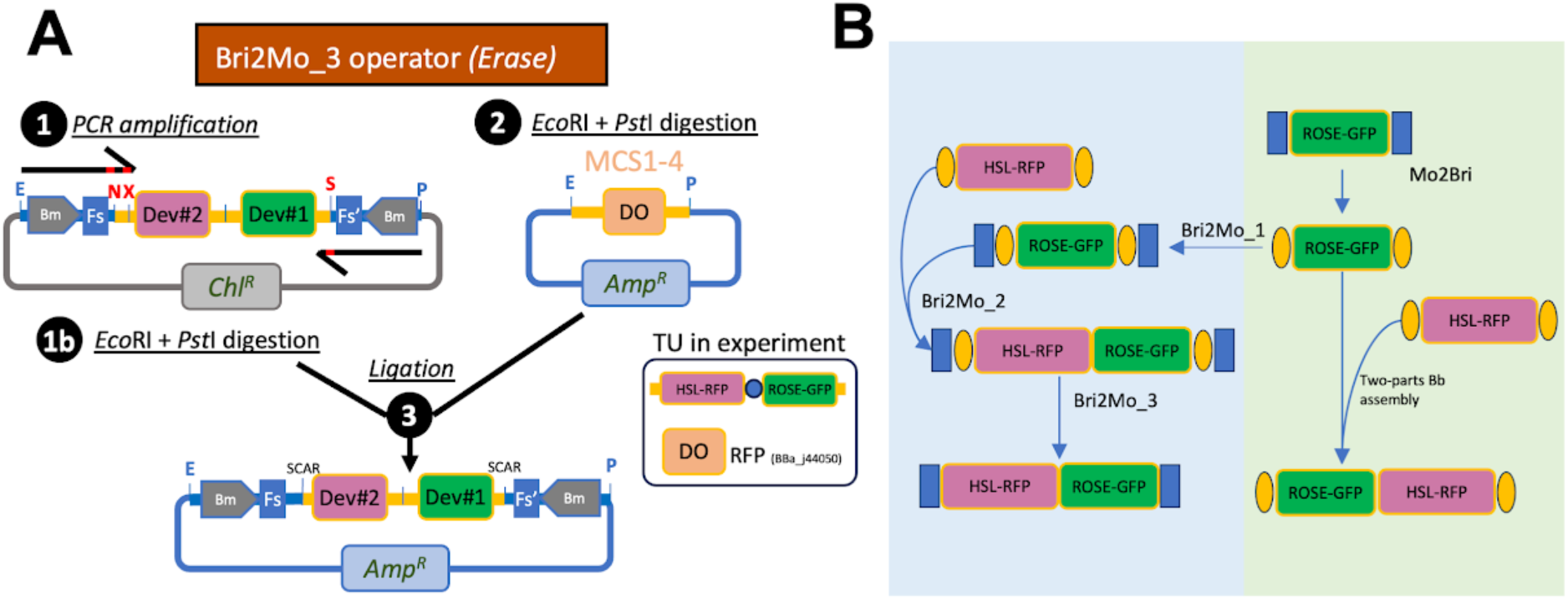
Implementation of the Bri2Mo_3 operator for the ablation of BioBrick recognition sequences (A). The workflow involves a site-directed mutagenic PCR (Step 1) employing the universal primer set: *Primer eraser forward* and *Primer eraser reverse* (refer to Supplementary Table S2). These oligonucleotides are designed with sequence-specific mismatches to eliminate the *Xba*I, *Not*I, and *Spe*I motifs while maintaining the structural integrity of the *BsmB*I, *EcoR*I, and *Pst*I sites. Sequential digestion of the PCR product and acceptor vector (Steps 1b and 2) allows for integration into any proposed SynBridge-compatible MCS. This procedure results in a construct containing exclusively MoClo Level 1 motifs. For experimental verification, MCS2 within pSB1K3 was utilized. Note that while diagrams illustrate backbones using chloramphenicol and ampicillin markers, antibiotic selection remains at the researcher’s discretion provided the donor and acceptor backbones utilize orthogonal resistances. **(B)** Comprehensive schematic of the validated conversion pathways established for the proof-of-concept validation in this study.

### Additional functionality of MCS2 for hierarchical MoClo integration

As it was previously anticipated, the architectural design of MCS2 incorporates *Bsa*I recognition sequences (marked with * in Figure 1D), enabling the plasmid to function directly as a MoClo Level 1 acceptor vector. During the *Bsa*I-mediated Golden Gate assembly, the legacy BioBrick restriction sites are inherently eliminated from the plasmid backbone. The resulting assembly yields a finalized transcriptional unit that is flanked exclusively by inward-facing *BsmB*I recognition targets. This arrangement prepares the assembled construct for subsequent higher-order MoClo iterations while ensuring the elimination of the original BioBrick sites.

Such a functionality was used in the assembly of the ROSE-GFP TU. In the case of HSL-RFP muti-TU device, TU1 (constitutive promoter - rbs - LuxR - terminator) and TU2 (Lux promoter - rbs – RFP - terminator) were independently assembled. Then, they were converted to a BioBricks format by a Mo2Bri operation to be joint in that standard. See details about the construction of these devices and for the DNA sequences in Supplementary Data.

### Functionality and interoperability

The combination of the Bri2Mo and Mo2Bri processes provides a set of formalized operations that enable interoperability between BioBrick and MoClo systems. As shown in Figure 1A, the entire procedure is complete in a closed loop workflow providing endless operation in an idempotent manner—both for the addition of new components and for the interconversion between the two systems—without any loss of functionality. The flowchart presents all possible steps and decision-making processes in an algorithmic manner, depending on the specific situation. As a proof of concept of a veritable interconversion within a close diagram loop Figure 4B summarize the interconversions of one to another assembly formats of ROSE-GFP and HSL-RFP, compatible with the idempotent addition of Biobricks parts.

## DISCUSSION

Historically, standardization has provided essential tools, rules, and best practices for both designing and characterizing parts (32). Working with standards offers a key advantage: a consensus framework that ensures component compatibility and interchangeability. However, the pursuit of better solutions leads to the emergence of alternatives, each shaped by distinct practical and historical factors. As a result, useful standard solutions coexist but are mutually incompatible, giving rise to conflicts that must be resolved through additional procedures.

Compared to traditional cloning, synthetic biology’s approach to DNA assembly exemplifies a hallmark of engineering: the pursuit of a generalized structured approach. Accordingly, this work addressed the interoperability between BioBricks and MoClo-like standards following the same philosophy. Our proposal introduces a methodology—a standard in itself—designed to maximize the use of genetic devices built with these mutually excluding assembly methods.

The integration of well characterized genetic devices is crucial for bottom-up process design (7). Assembling complex systems like metabolic pathways and regulatory circuits are examples of that. While current *de novo* synthesis remains limited by challenging sequences (e.g., high GC content or repetitive regions) (8), reusing existing constructs allows researchers to capitalize on previous efforts. Here, the ability to reversibly convert between MoClo and BioBricks offers complementary advantages: MoClo is highly efficient at Level 1 but becomes less flexible when scaling to higher-level assemblies, whereas BioBricks excels at joining multi-gene constructs in a pairwise (two-by-two) manner. By enabling a standardized conversion between both frameworks without ad-hoc adjustments, our system combines the strengths of both standards..This is exemplified by the bicistronic HSL-RFP construct in this work, in which individual TUs were assembled via MoClo and subsequently fused using BioBrick assembly. Interestingly, this construct represents a realistic scenario where PCR-based modification of flanking sites is unfeasible, as both TUs share identical terminators—the very regions typically targeted by tailored PCR strategies.

In an increasingly automated field (33) standardized conversions open the opportunity for the design of automated processes, a situation that is compatible with the demanding workflows operating in biofoundries (34). Interestingly, SynBridge offers an automatable solution to work with physical DNA collections of different precedence eluding the limitations of abovementioned traditional approximations based on site-directed mutagenesis.

A similar restriction applies to our methodology, specifically during the Bri2Mo_3 operation (the erasure of BioBrick scars). This limitation can be bypassed via *Xba*I and *Spe*I digestion, followed by DNA polymerase-mediated blunting and subsequent ligation. However, because this alternative relies on blunt-end ligation, assembly efficiency may be compromised, warranting further optimization.

SynBridge is revealed to be useful for the integration of higher-level functional units, such as transcriptional units, rather than the assembly or fusion of basic genetic parts (e.g., promoters, RBSs, etc.). While the conversion of simple parts is theoretically possible, it inevitably alters certain signals essential for proper function. This is the case of the distance between RBS and the start codon. This specific scope is reflected in our experimental validation, which focused exclusively on rescuing complete functional units.

Another aspect to mention is that our solution is easily transferred to any vector capable of accommodating our multiple cloning sites (MCS1-4). Although validated experimentally using pSB1C3, pSB1A3, and pSB1K3 backbones, Synbridge MCS are compatible with other established vectors, such as JUMP, pSB1C5, or SEVA 3.1. To enhance interoperativity, Figure S1 in Supplementary Data describes an additional MSC to directly convert the canonical SEVA 3.0/4.0 (35, 36) to a SEVA plasmid with *EcoR*I and *Pst*I flanking the cargo as it happens in SEVA 3.1 sibling (25) giving rise to a SynBridge-compatible backbone ready to use. These backbone formats work with SynBridge as they have been designed to support BioBrick standards. The auxiliary MCS5 described in Figure S1 in Supplementary Data allows such a modification by a double digestion *Pac*I/*Pst*I. According to this modification, all of these compatible backbones containing MCS1-5 preserve unique *EcoR*I and *Pst*I sites located at their MCS, a criterion to define SynBridge general architecture as described in Figure 1C.

Compatibility across all MoClo assembly steps requires vectors lacking internal *Bsa*I and *BsmB*I recognition sites. However, this is a condition not fully met by all standard BioBrick or SEVA plasmids configurations. For instance pSB1C3 and pSB1K3 contain from its origin an internal *BsmB*I chloramphenicol and kanamicyn resistance gene, respectively and pSB1A3 natively contains a *Bsa*I site. By its hand, for example SEVA181 and SEVA281 do not contain any of these two restriction sites but SEVA321 does it three times for *BsmB*I in its ampicillin resistance gene. Although certain SEVA vectors lack the *Bsa*I and *BsmB*I Type IIS recognition sites relevant to this study, ensuring full SynBridge compatibility is not entirely straightforward. They require verification that the modular parts used in a given SEVA configuration lack these problematic sites. However, in order to validate our methodology with the enzymes involved in our system we opted for the synbridge implementation pSB1C as *Bsa*I/*BsmB*I conflicts can be easily overcome by selecting the appropriate variant backbone, i.e. pSB1A3 for*BsmB*I involved processes and pSB1K3 and pSB1C3 when *Bsa*I is involved. In any case, digestion creates specific fusion sites of conflicting plasmids, however, after ligation (provided that there is no phosphatase activity) fusion sites should be reconstituted. If a MCS is required it can be easily transferred to a compatible plasmid via standard *EcoR*I and *Pst*I digestion. Interestingly, flanking the MCS with *EcoR*I and *Pst*I sites facilitates straightforward switching of antibiotic resistance markers, a desirable feature in cloning workflows.

Such a versatility of plasmid compatibility sums up to the idempotent condition maintained for additional integration of Biobricks pieces by Bri2Mo_2 operation. This operation can be iteratively applied until irreversible erasing of the biobricks sites is done. Concerning higher level iterations of MoClo, MCS1 and MCS2 are designed to host level 1 constructs. The MSC1 digestion with *BsmB*I allows it to accept level 1 constructs from most of the existent MoClo-like variants as they applied the definition of the same fusion site standardization for basic parts (15). This is the case for the abovementioned loop assembly, golden braid, golden and phytobricks among others. Conveniently for SynBridge, fusion sites have been quite conserved across different MoCLo-like assemblies. In this context, the use of a MoClo construct insert in MSC2 in other assembly systems is also possible. Digestion with *BsmB*I liberates DNA fragments compatible with the iterative assembly workflow of MoClo-like systems as they are liberated as the fusion of a level 1 of any of the Golden Gate standard assemblies referred before.

To conserve and expand this interoperativity at the level of backbones and assemblies our proposal inevitably introduces mandatory design principles for domesticated components: (1) MoClo devices should not contain internal restriction sites for BioBrick-associated enzymes. (2) BioBrick parts should ideally avoid *Bsa*I and other type IIS recognition sites (as detailed in Table 1). The use of accessory MCS (MSC3 and MSC4) described in Figure 1D to introduce new designed sequences can be benefited by avoiding *EcoR*V sites in the sequences of interest. Such a restriction maximizes the cloning options for the inclusion of these sequences to the SynBridge system by allowing a direct blunt end ligation from PCR amplification or DNA synthesis. Adhering to these part-domestication practices facilitates the bottom-up construction of complex DNA structures from curated libraries, establishing a repository free from interfering restriction sites.

Finally, experimental validation confirms that efficiency for cloning is comparable to those documented for Biobricks and MoClo systems. As expected, phenotypic behavior was not affected. This demonstrates that sequences from different standard sources can be effectively combined and reused, whether they originate from BioBricks or MoClo, within an endless closed loop of reversible conversion.

The results obtained optimizing the protocols reveal that efficiencies close to the 100% are possible under the tested conditions. However, such values might not be considered as a rule for any cloning. Applying these methodologies to larger constructs inevitably reduces efficiency, as plasmid stability and ligation processes are compromised when fragment size exceeds an optimal threshold.

Further research in cloning methodologies should focus on developing size-independent processes and streamlining one-pot reaction strategies. These advancements will significantly enhance our interchangeable framework, benefiting both manual and automated workflows. Ultimately, adhering to these design principles will ensure the long-term compatibility of genetic parts and facilitate seamless integration across standards.

In conclusion, SynBridge resolves a fundamental interoperability bottleneck between BioBricks and MoClo architectures. Through its bidirectional and idempotent pathways (Mo2Bri and Bri2Mo), the framework maintains complete sequence logic and functional dynamics across repeated standard conversions. By bridging legacy genetic parts with modern hierarchical assembly, SynBridge prevents part obsolescence, facilitates automated circuit construction, and provides a universal foundation to accelerate DBTL workflows in synthetic biology.

## SUPPLEMENTARY DATA

See Supplementary Data attached file.

## CONFLICT OF INTEREST

A patent application related to this work has been filed with the Spanish Patent and Trademark Office (OEPM) on June 8, 2026 Application No. P202630816, entitled ‘SynBridge: a versatile framework for bidirectional interconversion between BioBricks and MoClo assembly standards’.

## FUNDING

This work is supported by “II Plan Propio de la Universidad de Málaga (Ayuda E3) E30523”.

## AUTHOR CONTRIBUTIONS

CRC, AMC and MB conceived the idea and the experimental design. AMC and ANF conducted the experiments. AMC designed software development for image acquisition and automated phenotypic analysis. PMC conceived and constructed the ROSE-GFP unit, AMC and CRC analyzed the results. CRC, AMC, ANF and MB wrote and reviewed the manuscript.

## ACKNOWLEDGMENTS

The authors thank members of the B04-SIBIUMA group for their useful conversations and the anonymous reviewers for their valuable suggestions.

