## Supplementary Information for "SYNBRIDGE: A VERSATILE FRAMEWORK FOR BIDIRECTIONAL INTERCONVERSION BETWEEN BIOBRICKS AND GOLDEN GATE ASSEMBLY STANDARDS"

<sup>1</sup> Departamento Biología Molecular y Bioquímica, Universidad de Málaga, 29074, Málaga, Spain.

<sup>2</sup> Instituto de Investigación Biomédica de Málaga y Plataforma en Nanomedicina, IBIMA Plataforma BIONAND, 29590, Málaga, Spain

<sup>3</sup> Centro de Investigación en Red de Enfermedades Raras (CIBERER). Instituto de Salud Carlos III, 28029 Madrid, Spain.

#### 1. DNA parts and its sequences used in this work

**Explanation of Pseudosequence syntax:** To describe sequence architecture, a pseudo-sequence notation is used to represent the linear order of relevant genetic parts and restriction sites. Brackets ({ }) delineate previously described DNA sequences. For composite constructs, part names enclosed in brackets denote sequences that have been previously defined. Part definitions are presented in the table in hierarchical order, progressing from basic parts to composite devices. \*We adopt the concept of pseudosequence from the notion of pseudocode from software engineering: an informal, high-level description of an algorithm that mixes programming structure with human language.

**Table S1.** Sequences of all MCS shown in Figure 1. The sequence equivalent to the dropout, in this case RFP, is indicated with an underline. {DO=RFP} represents the transcriptional unit BBa\_J04450 used as a dropout gene for selection.

| MCS | Sequence |
| --- | --- |
| MCS1 Mo2Bri converter | 5'-TTAATTAAGAATTCGCGGCCGCTTCTAGAGGGAGTGAGACGGATATC(<u>DO=RFP</u>)GATATCCGTCTCACGCTTACTAGTAGCGGCCGCTGCAG -3' |
| MCS2 Bri2Mo converter | 5'-TTAATTAAGAATTCGGTCTCaGGAGGCGGCCGCGATATC(<u>DO=RFP</u>)GATATCACTAGTCGCTtGAGACCCTGCAG -3' |
| MCS3 BioBricks converter | 5'-TTAATTAAGAATTCGCGGCCGCTTCTAGAGGATATC(<u>DO=RFP</u>)GATATCTACTAGTAGCGGCCGCTGCAG -3' |
| MCS4 MoClo converter | 5'-TTAATTAAGAATTCCGTCTCaGGAGGATATC(<u>DO=RFP</u>)GATATCCGCTtGAGACGCTGCAGTCTAGA -3' |
| {DO=RFP} (RFP is Bba_J44050) | 5'-CAATACGCAAAACCGCCTCTCCCCGCGCTTGGCCGATTCAATATGCAGCTGGCACGACAGGTTTCCCGACTGGAAGCGGGCAGTGAGCGCAACGCAA TTAATGTGAGTTAGCTCACTCATTAGGCACCCAGGCTTTACACTTTATGCTTCCGGCTCGTATGTTGTGTGGAATTGTGAGCGGATAACAATTTACACATACT AGAGAAAGAGGAGAAATACTAGATGGCTTCTCCGAAGACGTTATCAAGAGTTTCATGCGTTTCAAAGTTTCGTATGGAAGGTTCCGTTAACGGTCACGAGTTT GAAATCGAAGGTGAAGGTGAAGGTGCGTCCGTACGAAGGTACCCAGACCGCTAAACTGAAAGTTACCAAAGGTGGTCCGCTGCCGTTTCGCTTGGGACATCCT GTCCCCGCAGTTCCAGTACGGTTCCAAAGCTTACGTTAAACACCCGGCTGACATCCCGGACTACCTGAAACTGTCTTCCCGGAAGGTTTCAAATGGGAACG TGTATGAACCTTCAAGACGGTGGTGTGTACCGTTACCCAGGACTCCTCCCTGCAAGACGGTGAGTTCACTACAAAGTTAAACTGCGTGGTACCAACTT CCGTCCGACGGTCCGGTTATGCAGAAAAAACCATGGGTTGGGAAGCTTCCACCGAACGTATGTACCCGGAAGACGGTGCTCTGAAAGGTGAAATCAAATG CGTCTGAAACTGAAAGACGGTGGCCACTACGACGCTGAAGTTAAACACCTACATGGCTAAAAAACCGGTTTCAGCTGCCGGGTGCTTACAAAACCGACATC AAACCTGGACATCACCTCCCAACGAAGACTACACCATCGTTGAACAGTACGAACGTGCTGAAAGGTGCTCACTCCACCGGTGCTTAATAACGCTGATAGTGT AGTGTAGATCGCTAGAGCCAGGCATCAATAAAACGAAAGGCTCAGTCAAGAACTGGGCTTTCGTTTATCTGTTGTTGTGCGGTGAACGCTCTCTAC TAGAGTCACACTGGCTCACCTTCGGGTGGGCTTTTCGCTTTATA_-3' |
| MCS 3.0 | 5'-TTAATTAAGAATTCGATATCCTGCAG -3' |
| MCS 3.0 conversion with DO | 5'-TTAATTAAGAATTCGATATC(<u>DO=RFP</u>)GATATCCTGCAG -3' |

**Table S2.** DNA sequence for primer eraser used by Bri2Mo<sub>3</sub> operation described in Figure 3 and in Figure 2S.

| Primer | Sequence |
| --- | --- |
| Primer eraser forward | 5'-GTATTGAATTCCGTCTCAGGAGTGAGACCGCGTCCGCTCCTAGAG-3' |
| Primer eraser reverse | 5'-GTAATCTGCAGCGTCTCAAGCGTGAGACCGCTAG-3' |

**Table S3.** DNA sequences and parts used in this work. To describe sequence architecture, a pseudo-sequence notation\* is used to represent the linear order of relevant genetic parts and restriction sites. Brackets ({} ) delineate previously described DNA sequences. For composite constructs, part names enclosed in brackets denote sequences that have been previously defined. Part definitions are presented in the table in hierarchical order, progressing from basic parts to composite devices. \*We adopt the concept of pseudosequence from the notion of pseudocode from software engineering: an informal, high-level description of an algorithm that mixes programming structure with human language.

| Part name | Sequence (Pseudosequence)* |
| --- | --- |
| Fs | 5'-GGAG-3' |
| Fs' | 5'-CGCT-3' |
| Biobricks prefix (long, according RCF-10) | 5'-GAATTCGCGGCCGCTTCTAGAG-3' |
| Biobricks suffix (according RCF-10) | 5'-TACTAGTAGCGGCCGCTGCAG-3' |
| Biobricks scar | 5'-TAGTAGAG-3' |
| BBa_J23100 (strong constitutive promoter) | 5'-TTGACGGCTAGCTCAGTCCTAGGTACAGTGCTAGC-3' |
| BBa_J23114 (weak constitutive promoter) | 5'-TTTATGGCTAGCTCAGTCCTAGGTACAATGCTAGC-3' |
| ROSE ( riboswitch, this study) | 5'-GCCGCGACAAGCGGTCCGGGCGCCCTAGGGGCCGCGCAGACGGGCGCCGAGGTGTCCGACGCCTGCTGGTTCCCATCTTGCTCCTTGAGAGGATTGGC-3' |
| BBa_B0030 (RBS medium) | 5'-ATTAAGAGGAGAAA-3' |
| BBa_E1010 (RFP) | 5'-ATGGCTTCCTCCGAAGACGTTATCAAAGAGTTCATGCGTTTCAAAGTTCGTATGGAAGGTTCCGTTAACGGTCACGAGTTCGAAATCGAAGGTGAAGGTGA AGGTCGTCCGTACGAAGGTACCCAGACCGCTAAACTGAAAGTTACCAAAGGTGGTCCGCTGCCGTTCGCTTGGGACATCCTGTCCCCGCAGTTCCAGTACG GTTCCAAAGCTTACGTTAAACACCCGGCTGACATCCCGGACTACCTGAAACTGTCCTTCCCGGAAGGTTTCAAATGGGAACGTGTTATGAACTTCGAAGACG GTGGTGTTGTTACCGTTACCCAGGACTCCTCCCTGCAAGACGGTGAGTTCATCTACAAAGTTAAACTGCGTGGTACCAACTTCCCGTCCGACGGTCCGGTTA TGCAGAAAAAAACCATGGGTTGGGAAGCTTCCACCGAACGTATGTACCCGGAAGACGGTGCTCTGAAAGGTGAAATCAAAATGCGTCTGAAACTGAAAGACG GTGGTCACTACGACGCTGAAGTTAAAACCACCTACATGGCTAAAAAACCGGTTCAGCTGCCGGGTGCTTACAAAACCGACATCAAACTGGACATCACCTCCC ACAACGAAGACTACACCATCGTTGAACAGTACGAACGTGCTGAAGGTCGTCACTCCACCGGTGCTTAATAACGCTGATAGTGCTAGTGTAGATCGC-3' |
| Bba_E4040 (GFP) | 5'-ATGCGTAAAGGAGAAGAACTTTTCACTGGAGTTGTCCCAATTCTTGTTGAATTAGATGGTGATGTTAATGGGCACAAATTTTCTGTCAGTGGAGAGGGTGAA GGTGATGCAACATACGGAAAACTTACCCTTAAATTTATTTGCACTACTGGAAAACTACCTGTTCCATGGCCAACACTTGTCACTACTTTCGGTTATGGTGTTCAA TGCTTTGCGAGATACCCAGATCATATGAAACAGCATGACTTTTTCAAGAGTGCCATGCCCGAAGGTTATGTACAGGAAAGAACTATATTTTTCAAAGATGACGG GAACTACAAGACACGTGCTGAAGTCAAGTTTGAAGGTGATACCCTTGTTAATAGAATCGAGTTAAAAGGTATTGATTTTAAAGAAGATGGAAACATTCTTGGACA CAAATTGGAATACAACTATAACTCACACAATGTATACATCATGGCAGACAAACAAAAGAATGGAATCAAAGTTAACTTCAAAATTAGACACAACATTGAAGATGGA AGCGTTCAACTAGCAGACCATTATCAACAAAATACTCCAATTGGCGATGGCCCTGTCCTTTTACCAGACAACCATTACCTGTCCACACAATCTGCCCTTTCGAA AGATCCCAACGAAAAGAGAGACCACATGGTCCTTCTTGAGTTTGTAACAGCTGCTGGGATTACACATGGCATGGATGAACTATACAAATAATAA-3' |
| Bba_C0062 (LuxR) | 5'-ATGAAAAACATAAATGCCGACGACACATACAGAATAATTAATAAAATTAAGCTTGTAGAAGCAATAATGATATTAATCAATGCTTATCTGATATGACTAAATGG TACATTGTGAATATTATTACTCGCGATCATTATCTCTATTCATGTTAAATCTGATATTTCAATCCTAGATAATTACCCTAATAAATGGAGGCAATATTATGATGA CGCTAATTTAATAAAATATGATCCTATAGTAGATTATTCTAACTCCAATCATCCCAATTAATTGGAATATATTTGAAAAACAATGCTGTAAATAAAAAATCTCCAAAT GTAATTAAAGAAGCGGAAAAACATCAGGCTTATCACTCGGGTTAGTTTCCCTATTTCATACGGCTAACAAATGGCCTCGGAATGCTTAGTTTTGCACATTGAGAAAAA GACAACTATATAGATAGTTTATTTTACATGCGGTGATGAACATACCATTAATTGTTCTCTCTAGTTGATAATTATCGAAAAATAAATATAGCAAAATAAATAATCAAA CAACGATTTAACCAAAAGAGAAAAAGAATGTTTAGCGTGGGCGATCGGAAGGAAAAAGCTCTTGGGATATTTCAAAAATATTAGGTTGCAGTGAGCGTACTGTCA CTTTCCATTTTAACCAATCGCGAAATGAAACTCAATGAAACAAACCCGCTGCGAAAGTATTTTCAAGAACTTTTAAACAGGAGCAATTTAATGCCCCTACTTTAAAAA TTAATAA-3' |
| BBa_R0062 (pLux) | 5'-ACCTGTAGGATCGTACAGGTTTACGCAAGAAAATGGTTTGTTATAGTCGAATAAA-3' |
| BBa_B0015 (Terminator) | 5'-CCAGGCATCAAATAAAACGAAAGGCTCAGTCGAAAGACTGGGCCTTTCGTTTTATCTGTTGTTTGTCGGTGAACGCTCTCTACTAGAGTCACACTGGCTCA CCTTCGGGTGGGCCTTTCTGCGTTTATA-3' |
| ROSE-GFP (TU MoClo assembled) | 5'-{Bba_J23100}-{ROSE}-{Bba_E4040 (GFP)}-{Bba_B0015}-3'<br>5'-{Fs}-TTGACGGCTAGCTCAGTCCTAGGTACAGTGCTAGCTACTGCCGCGACAAGCGGTCCGGGCGCCCTAGGGGCCCGGCGCAGACGGGCGCCGGAGG TGTCCGACGCGCTGCTGGTTCCTCATTTGCTCCTTGGAGGATTTGGCAATGCGTAAAGGAGAAGAACTTTTCACTGGAGTTGTCCCAATTCCTGTTGAATTAGA TGGTGATGTTAATGGGACAAATTTTCTGTCACTGGAGAGGGTGAAGGTGATGCAACATACGGGAAACTTACCCCTAAATTTATTTGCACACTGGAAGAACTAC CTGTTCCAGTGGCCAACTGTCACACTATTTTCGGTTATGGTGTTCATGCTTTGCGAGATACCCAGATCATATGAAACAGCAGTACGTTTTCAAGATGGCCATG CCGGAAGGTTATGTACAGGAAAGTATATTTCAAAGATGACGGGAACGCAACAGACGCTGCTGAAGTCAAGTTTGAAGGTGATACCTTGTTAATGAAT CGAGTTAAAGGTATTGATTTTAAAGAAGATGGAACATTCTTGGACACAATTTGGAATACAACATAACTCACACAATGTATACATCATGGCAGACAAAACAAAAG AATGGAATCAAAGTTAACTTCAAATTTAGACACAACATTGAAGATGGAAGCGTTCAACTAGCAGACCATTTATCAACAAAATCTCCAATTTGGCGATGGCCCTGTG CTTTTACACAGAACCATTACCTGTCCACACAATCTGCCCTTTCCGAAGATCCCAACGAAAAAGAGACCCACATGGTCTCTTGCAGATCTGTAAACAGCTGCTG GGATTACACATGGCATGGATGAACATATACAAATAAACAGGCATCAATAAAAAACGAAAGGCTCAGTCGAAAGACTGGGCCCTTCGTTTTATCTGTTGTTTGTG CCGTGAACGCTCTCTACTAGAGTCACACTGGCTCACCTTCGGGTGGGCCCTTCTGCGTTTATA-{Fs}-3' |
| HSL-RFP (Multi-TU, each TU assembled by MoClo, TU fusion by Biobricks) | 5'-{Biobricks prefix} - {Fs} - {BBa_J23114} - {Bba_B0010 (RBS)} - {Bba_C0062} - {Bba_B0015} - {Fs} - {Biobrick scar} - {Fs} - {BBa_R0062 (pLux)} - {Bba_B0010 (RBS)} - {Bba_E1010} - {Bba_B0015} - {Fs} - {Biobricks suffix} - 3'<br>5'-{Fs}-GGAGTTTATGGCTAGCTCAGTCCTAGGTACAATGCTAGCTACTATTAAAGAGGAGAAATACTAAATGAAAAACATAAATGCCGACGACACATACAGAATA ATTAATAAAATTAAAGCTTGTAGAAGCAATAATGATATTAATCAATGCTTATCTGATATGACTAAAATGGTACATTGTGAATATTATTTACTCGCGATCATTTATCCTCA TTCTATGGTTAAATCTGATATTTCAATCCTAGATAATTACCCTAAAAAATGGAGGCAATATTATGATGACGCTAATTTAATAAAATATGATCCTATAGTAGATTATTCTA ACTCCAATCATTCACCAATTAATTGGAATATATTTGAAAACAATGCTGTAAATAAAAAATCTCCAAATGTAATTAAAGAAGCGAAAACATCAGGTCTTATCACTGGG TTTAGTTTCCCTATTCATACGGCTAACAATGGCTTCGGAATGCTTAGTTTTGCACATTCAGAAAAAGACAACTATATAGATAGTTTATTTTTACATGCGTGTATGAA CATACCATTAATTGTTCCTTCTCTAGTTGATAATTATCGAAAAATAAATATAGCAAATAATAAATCAAACAACGATTTAACCAAAAGAGAAAAAGAATGTTTAGCGTG GGCATGCGAAGGAAAAAGCTCTTGGGATATTTCAAAAATATTAGGTTGCAGTGAGCGTACTGTCACTTTCCATTTAACCAATGCGCAAATGAAACTCAATACAAC AAACCGCTGCCAAAGTATTTCTAAAGCAATTTTAACAGGAGCAATTGATTGCCCATACTTTAAAAATTAATAAGCTTCCAGGCATCAAATAAAACGAAAGGCTCA |

GTCGAAAGACTGGGCCCTTTCGTTTATCTGTTGTTGTGCGGTGAACGCTCTCTACTAGAGTCACACTGGCTCACCTTCGGGTGGGCCCTTCTGCGTTTATACG
CTTACTAGAGGGAGACCTGTAGGATCGTACAGGTTTACGCAAGAAATGGTTTGTATAGTCGAATAAATACTATTAAAGAGGAGAAATACTAAATGGCTTCCTC
CGAAGACGTTATCAAAGAGTTCATGCGTTTCAAAGTTCGATGGAAGGTTCCGTTAACCGGTCACGAGTTCGAAATCGAAGGTCAAAGGTCGTCGCTG
CGAAGGTACCCAGACCGCTAAACTGAAAGTTACCAAAGGTGGTCCGCTGCCGTTCCGTTGGGACATCCTGTCCCCGACGTTCAGTACGGTTCCAAAGCTTA
CGTTAAACACCCCGCTGACATCCCGGACTACCTGAAACTGTCCTTCCCGGAAGGTTTCAAATGGGAACGTTGTTAGAACTTCGAAGACCGGTGGTGTGTTAC
CGTTACCACGAGCTCTCCCTGCAAGACGGTGAGTTCATCTACAAAGTTAAAGTCGGTGGTACCAACTTCCCGTCGAGCGTCGGGTTATGCAGAAAAAAC
CATGGGTTGGGAAGCTTCCACCGAACGTATGTACCCGGAAGACGGTGCTCTGAAAGGTGAAATCAAATCGCTGTGAAACTGAAAGACCGGTGGTCACTACGAA
CGCTGAAGTTAAAACCACCTACATGGCTAAAAAACCGGTTCAGCTGCCGGGTGCTTACAAAACCGACATCAAACTGGACATCACCTCCCACAACGAAGACTA
CACCATCGTTGAACAGTACGAACGTGCTGAAGGTCGTCACTCCACCGGTGCTTAATAACGCTGATAGTGCTAGTGTAGATCGCGCTTCCAGGCATCAAATAAA
ACGAAAGGCTCAGTCGAAAGACTGGGCCTTTCGTTTTATCTGTTGTTTGTCGGTGAACGCTCTCTACTAGAGTCACACTGGCTCACCTTCGGGTGGGCCTTT
TCGCGTTTATA-[Fs]-3'

pSB1C3
(BioBrick Backbone)

TACTAGTAGCGGCGCTGCAGTCCGGCAAAAAAGGGCAAGGTGTACCAACCCGTGCCCTTTTTCTTTAAACCGAAAAAGATTACTTCGCGTTATGCAGGCTTCC
TCGCTCACTGACTCGCTGCGCTCGGTCTGTTGCGCTGCGGCGAGCGGTATCAGCTCACTCAAAGGCGGTAAATACGGTTATCCACAGAATCAGGGGATAACGC
AGGAAAAAACAATGTGAGCAAAAGGCCAGCAAAAGGCCAGGAACCGTAAAAAGGCCGCGTTGCTGGCGTTTTTCCACAGGCTCCGCCCCCTGACGAGCAT
CACAAAAATCGACGCTCAAAGTCAGAGGTGGCGAAACCCGACAGGACTATAAAGATACCAAGCGCTTCCCGCTGGAAGCTCCCTCGTCCGCTCTCTGTTCC
GACCCTGCCGCTTACCGGATACCTGTCCGCCCTTCTCCCTTCGGGAAGCGTGGCGCTTTCTCATAGCTACAGCTGTAGGTATCTCAGTTCGGGTGATAGTCTGT
TCGCTCCAAGCTGGGCTGTGTGCACGAACCCCGCTTCAGCCCGACCGCTGCGCCTTATCCGGTAACTATCGTCTTGAGTCCAACCCCGTAAAGACACGACT
TATCGCCACTGGGACAGCCACTGGTAACAGGATTAGCAGAGCGAGGTATGAGCGGTGCTACAGAGTTCTTGAAGTGGTGAGCTAACCTACCGCTACACTA
GAAGAACAGTATTTGGTATCTGCGCTCTGCTGAAGCCAGTTACCTTCGGGAAGAAAGATTTGGTAGCTCTTGATCCGGCAACAAACCCAGCGGTGGTAGCGGTG
GTTTTTTTGTGTCGAAGCAGCAGATTACGCGCAGAAAAAAGGATCTCAAGAAGATCCTTTGATCTTTTCTACGGGCTGTAGCTGTGGAACGAAAACTC
ACGTTAAGGGATTTTGGTCATGAGATTATCAAAAAGGATCTTCACTAGATCCTTTTAAATTAATAAGTTTTAAATCAATCTAAAGTATATATGAGTAACTTG
GTCTGACAGCTCGAGGCTTGGATTCTCACCATAAAAAACGCCGCGGCCAACCCGAGCGGTTCTGAACAAATCCAGATGGAGTTCTGAGGTCACTTACCTGGATC
TATCAACAGGAGTCCGAAGCGAGCTGATATCAAATTACGCCCGCCCTGCCACTCATCGCAGTACTGTTGTAATTCATTAAGCATCTGCCGACATGGAAGCCA
TCACAAACGGCATGATGAACCTGAATCGCCAGCGGCATCAGCACCTTGTGCGCTTGCGTATAATATTGCCCATGGTGAAACCGGGGCGGAAGAGTTGTCC
ATATTGGCCACGTTTAAATCAAACTGGTGAACCTCACCAGGGATTTGGCTGAGCAGCAAAAAATATTCTCAATAAACCCTTTAGGGAATAAGCCAGGTTTTTC
ACCGTAACAGCCCACTTTTCCGAATATGTGTAGAAACTGCCGGAATGCGTGTTACTCACTCCAGAGCGATGAAGACGTTTCAGTTTGTCTATGGAAAAA
CGGTGTAAACAGGGTGAACACTATCCCATATCACCAGCTCACCCTCTTCATTGCGCATACGAAATCCGGATAGGACATTATCAGCGCGGCAAGAATGTGAATA
AAGCGCCGATAAAACTTGTGCTTATTTTCTTACGGCTTTTAAAGGCCGTATATCAGAGTAAATAAATCCTTAGCTTTTCGTAAGATGATTTCGGAATTCGCCGCCCTTC
GCCTCAAAATGTTCTTTACGATGCCATTGGGATATATCAACGGTGGTATATCCAGTGATTTTTTTCTCCATTTTAGCTTCCTTAGCTCCTGAAAATCTCGATAACTC
AAAAAATACGCCCGGTAGTGATCTTATTTCATTATGGTGAAAGTTGGAACCTCTTACGTGCCCGATCAACTCGAGTGCCACCTGACGTCTAAGAAACCATTATTA
TCATGACATTAACCTATAAAAATAGGCGTATCACGAGGCAGAATTTCAGATAAAAAAAATCCTTAGCTTTCGCTAAGGATGATTTCTGGAATTCGCGGCCGCTTC
TAGAG

pSB1K3 (BioBrick
Backbone)

TACTAGTAGCGGCCGCTGCAGTCCGGCAAAAAAGGGCAAGGTGTCACCACCCTGCCCTTTTTCTTTAAAACCGAAAAGATTACTTCGCGTTATGCAGGCTTCC
TCGCTCACTGACTCGCTGCGCTCGGTCGTTCGGCTGCGGCGAGCGGTATCAGCTCACTCAAAGGCGGTAATACGGTTATCCACAGAATCAGGGGATAACGC
AGGAAAGAACATGTGAGCAAAAGGCCAGCAAAAGGCCAGGAACCGTAAAAAGGCCGCGTTGCTGGCGTTTTTCCACAGGCTCCGCCCCCCTGACGAGCAT
CACAAAAATCGACGCTCAAGTCAGAGGTGGCGAAACCCGACAGGACTATAAAGATACCAGGCGTTTCCCCCTGGAAGCTCCCTCGTGCGCTCTCCTGTTCC
GACCCTGCCGCTTACCGGATACCTGTCCGCCTTTCTCCCTTCGGGAAGCGTGGCGCTTTCTCATAGCTCACGCTGTAGGTATCTCAGTTCGGTGTAGGTCGT
TCGCTCCAAGCTGGGCTGTGTGCACGAACCCCCCGTTCAGCCCGACCGCTGCGCCTTATCCGGTAACTATCGTCTTGAGTCCAACCCGGTAAGACACGACT
TATCGCCACTGGCAGCAGCCACTGGTAACAGGATTAGCAGAGCGAGGTATGTAGGCGGTGCTACAGAGTTCTTGAAGTGGTGGCCTAACTACGGCTACACTA
GAAGAACAGTATTTGGTATCTGCGCTCTGCTGAAGCCAGTTACCTTCGGAAAAAGAGTTGGTAGCTCTTGATCCGGCAAACAAACCACCGCTGGTAGCGGTG
GTTTTTTTGTTTGCAAGCAGCAGATTACGCGCAGAAAAAAAGGATCTCAAGAAGATCCTTTGATCTTTTCTACGGGGTCTGACGCTCAGTGGAACGAAAACTC
ACGTTAAGGGATTTTGGTCATGAGATTATCAAAAAGGATCTTCACCTAGATCCTTTTAAATTAAAAATGAAGTTTTAAATCAATCTAAAGTATATATGAGTAAACTTG
GTCTGACAGCTCGAGTCCCGTCAAGTCAGCGTAATGCTCTGCCAGTGTTACAACCAATTAACCAATTCTGATTAGAAAAACTCATCGAGCATCAAATGAAACTG
CAATTTATTCATATCAGGATTATCAATACCATATTTTTGAAAAAGCCGTTTCTGTAATGAAGGAGAAAACTCACCGAGGCAGTTCCATAGGATGGCAAGATCCTG
GTATCGGTCTGCGATTCCGACTCGTCCAACATCAATACAACCTATTAATTTCCCCTCGTCAAAAATAAGGTTATCAAGTGAGAAATCACCATGAGTGACGACTGA
ATCCGGTGAGAATGGCAAAAGCTTATGCATTTCTTTCCAGACTTGTTCAACAGGCCAGCCATTACGCTCGTCATCAAAATCACTCGCATCAACCAAACCGTTAT
TCATTCGTGATTGCGCCTGAGCGAGACGAAATACGCGATCGCTGTTAAAAGGACAATTACAAACAGGAATCGAATGCAACCGGCGCAGGAACACTGCCAGCG
CATCAACAATATTTTCACCTGAATCAGGATATTCTTCTAATACCTGGAATGCTGTTTTCCCGGGGATCGCAGTGGTGAGTAACCATGCATCATCAGGAGTACGGA
TAAAATGCTTGATGGTCGGAAGAGGCATAAATTCCGTCAGCCAGTTTAGTCTGACCATCTCATCTGTAACATCATTGGCAACGCTACCTTTGCCATGTTTCAGA
AACAACTCTGGCGCATCGGGCTTCCCATACAATCGATAGATTGTCGCACCTGATTGCCCGACATTATCGCGAGCCCATTTATACCCATATAAATCAGCATCCATG
TTGGAATTTAATCGCGGCTGGAGCAAGACGTTTCCCGTTGAATATGGCTATAACACCCCTTGTATTACTGTTTATGTAAGCAGCAAGCTTTTATTGTTTCATGAT
GATATATTTTATCTTGTGCAATGTAAACATCAGAGATTTTGAGACACAACCGTGGCTTTGTTGAATAAATCGAACTTTTGTCTGAGTTGAAGGATCAGCTCGAGTGC
CACCTGACGCTTAAGAAACCATATTATCATGACATTAACTTATAAAAATAGGCGTATCAGGAGCAGAATTTTCAGATAAAAAAATCCTTAGCTTTTCGTAAGGA
TGATTTCGGAATTCGCGGCCGCTTCTAGAG

pSB1A3 (BioBrick
Backbone)

TACTAGTAGCGGCCGCTGCAGTCCGGCAAAAAAGGGCAAGGTGTCACCACCCTGCCCTTTTTCTTTAAAACCGAAAAGATTACTTCGCGTTATGCAGGCTTCC
TCGCTCACTGACTCGCTGCGCTCGGTCGTTCGGCTGCGGCGAGCGGTATCAGCTCACTCAAAGGCGGTAATACGGTTATCCACAGAATCAGGGGATAACGC
AGGAAAGAACATGTGAGCAAAAGGCCAGCAAAAGGCCAGGAACCGTAAAAAGGCCGCGTTGCTGGCGTTTTTCCACAGGCTCCGCCCCCCTGACGAGCAT
CACAAAAATCGACGCTCAAGTCAGAGGTGGCGAAACCCGACAGGACTATAAAGATACCAGGCGTTTCCCCCTGGAAGCTCCCTCGTGCGCTCTCCTGTTCC
GACCCTGCCGCTTACCGGATACCTGTCCGCCTTTCTCCCTTCGGGAAGCGTGGCGCTTTCTCATAGCTCACGCTGTAGGTATCTCAGTTCGGTGTAGGTCGT
TCGCTCCAAGCTGGGCTGTGTGCACGAACCCCCCGTTCAGCCCGACCGCTGCGCCTTATCCGGTAACTATCGTCTTGAGTCCAACCCGGTAAGACACGACT
TATCGCCACTGGCAGCAGCCACTGGTAACAGGATTAGCAGAGCGAGGTATGTAGGCGGTGCTACAGAGTTCTTGAAGTGGTGGCCTAACTACGGCTACACTA
GAAGAACAGTATTTGGTATCTGCGCTCTGCTGAAGCCAGTTACCTTCGGAAAAAGAGTTGGTAGCTCTTGATCCGGCAAACAAACCACCGCTGGTAGCGGTG
GTTTTTTTGTTTGCAAGCAGCAGATTACGCGCAGAAAAAAAGGATCTCAAGAAGATCCTTTGATCTTTTCTACGGGGTCTGACGCTCAGTGGAACGAAAACTC
ACGTTAAGGGATTTTGGTCATGAGATTATCAAAAAGGATCTTCACCTAGATCCTTTTAAATTAAAAATGAAGTTTTAAATCAATCTAAAGTATATATGAGTAAACTTG
TTGGAATTTAATCGCGGCTGGAGCAAGACGTTTCCCGTTGAATATGGCTATAACACCCCTTGTATTACTGTTTATGTAAGCAGCAAGCTTTTATTGTTTCATGAT
GATATATTTTATCTTGTGCAATGTAAACATCAGAGATTTTGAGACACAACCGTGGCTTTGTTGAATAAATCGAACTTTTGTCTGAGTTGAAGGATCAGCTCGAGTGC
CACCTGACGCTTAAGAAACCATATTATCATGACATTAACTTATAAAAATAGGCGTATCAGGAGCAGAATTTTCAGATAAAAAAATCCTTAGCTTTTCGTAAGGA
TGATTTCGGAATTCGCGGCCGCTTCTAGAG

ROSE-GFP in the
donor backbone
(MCS2 cut with BsaI
from level 1 TU
assembly)

5'-[EcoRI]-[BsmBI]-[Fs]-[ROSE-GFP]-[Fs']-[-BsmBI]-[PstI]-3'
5'-GAATTCGCTCTCAGGAGTTGACGGCTAGCTCAGTCTAGGTCAGTACTGCTGCGCGACAAGCGGTCCGGGCGCCCTAGGGGCGCCGGCGCAGA
CGGGCGCGGAGGTGTGCGACGCCGTGCTGGTCCCATCTTGTCTCTTGGAGGATTTGGCAATGCGTAAAGGAGAAGAACTTTTCTAGTGGAGTTGTCCCAAT
TCTTGTGAATAGATGGTGATGTTAATGGGCACAAATTTTCTGTCAGTGGAGGAGGTTGAAGGTGATGCAACATACCGGAAAACTTACCCTTAAATTTATTTGCAC
TCTGTTGCAATCGGCTTTCATGGCCAACTGTTCACTACTTTTCGGTTATGGTGTTCATGCTTTTGGGAGATACCCAGTATGAAACAGCATGATGTTTTC
CAAGAGGTGCCATCGCCGAAGGTTATGTACAGGAAAGAACTATATTTTCAAAGATGACGGGAACTACAAGACACGTGCTGAAGTCAAGTTTGAAGGATGATACC
CTTGTTAATGAGTCCGTTAAAGGTATTGATTTTAAAGAGATGGAACACTCTTGGACACAAATTTGGAATACAACTATGACTACACAACTGATACATGAGCGG
AGACAAACAAAGAAATGGAATCAAGTTAACTTCAAATATAGACACAACATTGAAGATGGAAGCGTTCAACTAGCAGACCATATCAACAAATATCTCAAATTTGG
CGATGGCCCTGTCTTTTACCAGACAACCATTACCTGTCCACACAATCTGCCCTTTGCAAGATGCCAACGAAAGGAGAGACCACTGGTCCCTCTTGTAGTTT
GTAACGCTGCTGGGATTACACATGGCATGGATGAACATATAAATAACCAAGGCATCAATCAAAATAACGAAAGGATGAGCTGCAAGAGCTGGGCGCTTTTCTGTTT
ATCTGTTGTTGTGCGGTGAACGCTCTCTACTAGAGTCACACTGGCTCACCTTCGGGTGGGCCCTTTCGCGTTTATACGCTTGAGACGCTGCAG-3'

HSL-RFP within
MCS1

5'-[EcoRI]-[NotI]-[XbaI]-[Fs]-[HSL-RFP]-[Fs']-[SpeI]-[NotI]-[PstI]-3'
5'-GAATTCGCGGCGCGCTTCTAGAGGGAGGGAGTTTATGGCTAGCTCAGTCTAGGTACAATGCTAGCTACTATTAAGAGGAGAGAAATACTAAATGAAAAACATAA
ATGCCGACGACACATACAGAATAATTAATAAAATTAAAGCTTGTAGAAGCAATAATGATATTAATCAATGCTTATCTGATATGACTAAAATGGTACATTGTGAATATTA
TTTACTCGCGATCATTTATCCTCATTCTATGGTTAAATCTGATATTTCAATCCTAGATAATTACCCTAAAAAATGGAGGCAATATTATGATGACGCTAATTTAATAAAA
TATGATCCTATAGTATGATTATCTTAACCTCAACATTAACCAATTAATTTGGAATATATTTGAAAACAATGCTGTAAATAAAAAATCTCAAATGTAAATTAAGAACGGA
AAACACTAGGCTTTATCACTGGGTTAGTTTTCCTTATCATCGGCTAAACAATGGCTTCCGGAATGCTTAGTTTTCACATTAGAAAAAGCAACTATATAGATGATG
TTTATTTTACATGCGGTGATGAACATACCAATTAATTTGTTCTTCTAGTTGATAATTATCGAAAAATAAATATAGCAAAATAAATAACCAACGATTAAACCAAA

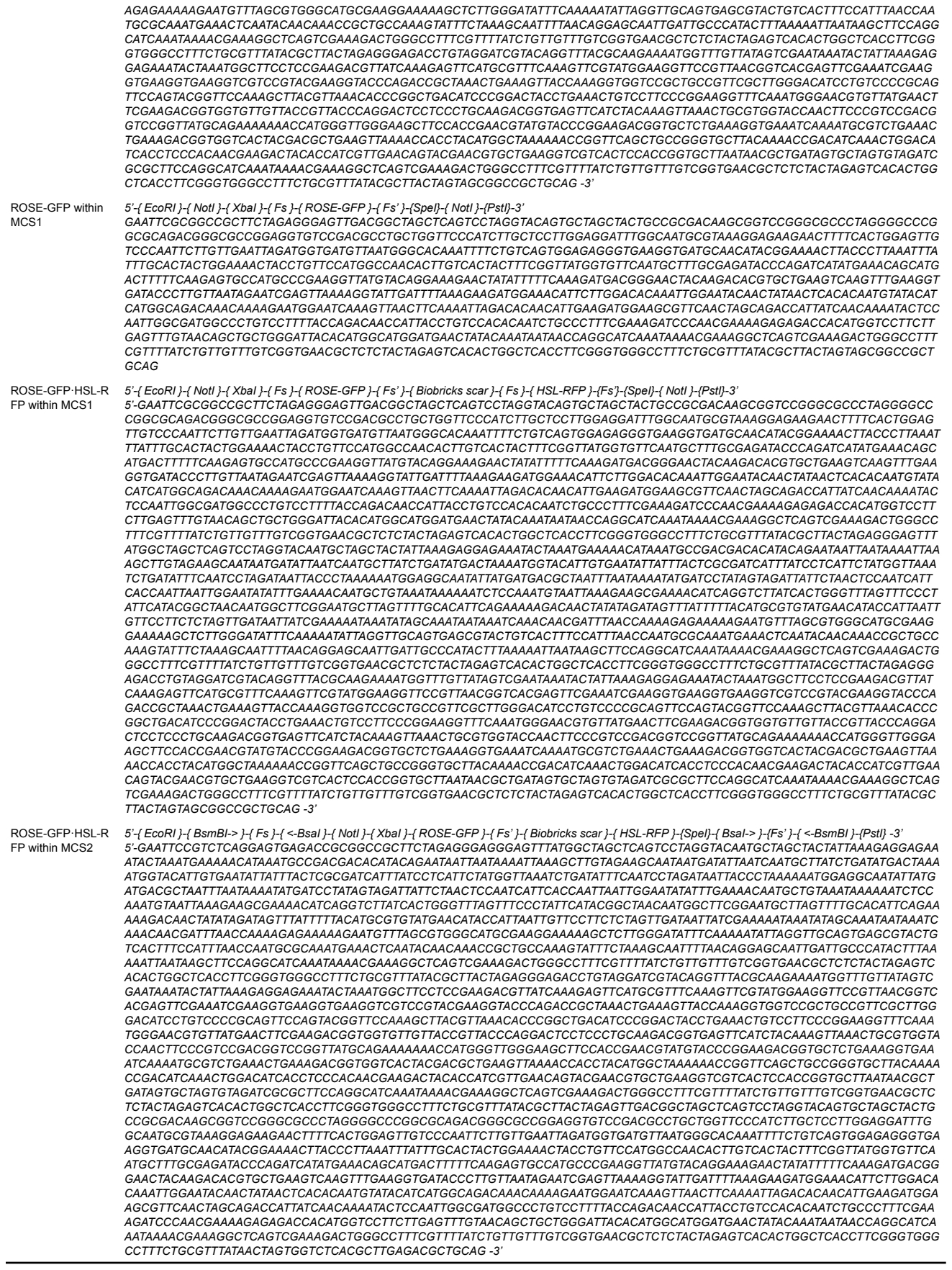

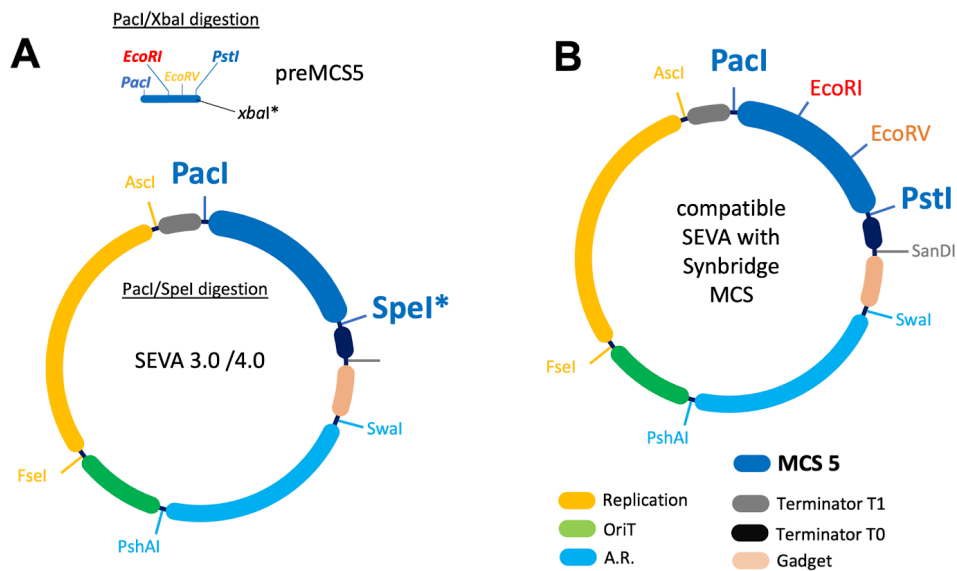

**Figure S1.**

**Figure S1. Auxiliary MCS5 generation.** BioBrick-compatible SEVA systems can be generated from SEVA 3.0 or SEVA 4.0 vectors. Starting from a SEVA Cargo 1 plasmid, these systems are constructed by cloning the PacI/XbaI-digested synthetic preMCS5 fragment (B) into PacI/SpeI-digested pSEVA backbones, exploiting the compatible cohesive ends generated by XbaI and SpeI. Note that the EcoRI/PstI restriction site pair enables its function as a BioBrick backbone. Additionally, an integrated EcoRV site permits the insertion of blunt-ended DNA fragments derived from gene synthesis or PCR amplification.

#### 2. Implementations of departing constructs for experimental validation.

##### ROSE-GFP TU

###### *ROSE domestication.*

Basic part sequence were obtained from BBa\_K115001 from the Part Registry https://registry.igem.org/parts/BBa_K115001. This sequence was originally taken from the *Bradi rhizobium Japonicum* as the 5'UTR (ROSE) of a heat shock protein (prfA). The sequence was described by (1) The analysis of the sequence showed that its sequence was incompatible with Golden Gate. Consequently, the native ROSE sequence was retrieved, and nucleotide substitutions A95>T and T123>A were introduced to ensure Golden Gate compatibility while preserving its secondary structure.

###### *GFP-ROSE construction.*

GFP-ROSE was designed to have a constitutive promoter, the ROSE riboregulator, a GFP and terminator. The specific parts were: BbaJ23100+ROSE+Bba\_E4040(GFP)+Bba\_B0015. The construct came from the Bsal-Level.

The acceptor plasmid is a pSB1C3 with MCS2 due to its functionality of being an acceptor plasmid for Level 0 assemblies when digested with *Bsa*I. ROSE-GFP within MCS2 after modular cloning has the following representation in a pseudosequence format.

5'-{EcoRI} - {BsmBI} - {Fs} - {ROSE-GFP}{Fs'} - {<BsmBI}-{PstI}-3' See Table S1 for details of the sequence.

Fs and Fs' coincide with 5'- fusion site promoter and 3' fusion site terminator of the TU. BsmBI recognition sites are oriented inwards, to the fusion sites, giving rise to a canonical Level1 structure of a MoClo.

#### HSL-RFP

HSL-RFP is a bicistronic device built in the following manner:

##### *1. Level 1 MoClo assembly:*

TU1: Lux promoter with a rbs followed by an RFP and terminator, more precisely:

{EcoRI} - {BsmBI} - {Fs} - {BBa\_J23114} - {Bba\_B0010 (RBS)} - {Bba\_C0062} - {Bba\_B0015} - {Fs'} - {<BsmBI}-3'.

This assembly was done in a pSB1A3 with MCS2 within *EcoRI/PstI* sites that was cut by *Bsa*I acting as the acceptor backbone of a Level 1 MoClo assembly.

TU2: {Fs} - {EcoRI} - {BsmBI} - {BBa\_R0062 (pLux)} - {Bba\_B0010 (RBS)} - {Bba\_E1010} - {Bba\_B0015} - {Fs'} - {<BsmBI}-3'

This assembly was done in a pSB1A3 with MCS2 within *EcoRI/PstI* sites that it was cut by *Bsa*I acting as acceptor backbone of a Level 1 MoClo assembly.

##### *2. Conversion to BioBricks using Mo2Bri operator:*

We obtained a

TU1:

5'-{BioBricks prefix} - {Fs} - {BBa\_J23114} - {Bba\_B0010 (RBS)} - {Bba\_C0062} - {Bba\_B0015} - {Fs'} - {BioBricks suffix}-3'

TU2:

5'-{BioBricks prefix} - {Fs} - {BBa\_R0062 (pLux)} - {Bba\_B0010 (RBS)} - {Bba\_E1010} - {Bba\_B0015} - {Fs'} - {BioBricks suffix}-3'

##### *3. A Canonical BioBricks assembly using as a backbone acceptor pSB1C3:*

The result was a constitutive production of LuxR followed by regulatable production of RFP by pLux promoter. More precisely in a pseudo-sequence format:

5' -{BioBricks prefix} - {Fs} - {BBa\_J23114} - {Bba\_B0010 (RBS)} - {Bba\_C0062} - {Bba\_B0015} - {Fs'} - {BioBrick scar} - {Fs} - {BBa\_R0062 (pLux)} - {Bba\_B0010 (RBS)} - {Bba\_E1010} - {Bba\_B0015} - {Fs'} - {BioBricks suffix}-3'

### Synbridge protocols

The following tables describe the protocols related to Synbridge

#### Time incubation protocols:

**Table S4. SynBridge time incubation protocols.**

| Operation | Protocol |
| --- | --- |
| Mo2Bri one-pot (42°C) | 29 cycles:<br>Digestion: 42 °C (2 min)<br>Ligation: 16 °C (5 min)<br>Digestion: 42 °C (5 min) → 60 °C (5 min)<br>Digestion: 42 °C (5 min) → 80 °C (5 min) |
| Mo2Bri one-pot (37°C) | 29 cycles:<br>Digestion: 37 °C (2 min)<br>Ligation: 16 °C (5 min)<br>Digestion: 37 °C (5 min) → 60 °C (5 min)<br>Digestion: 37 °C (5 min) → 80 °C (5 min) |
| Mo2Bri sequential (rSAP+) | Digestion: 37 °C (20 min) → 80 °C (20 min)<br>Dephosphorylation of backbone DNA: 37 °C (30 min) → 80 °C (5 min)<br>Ligation: 25 °C (30 min) → 65 °C (10 min) |
| Mo2Bri sequential (rSAP-) | Digestion: 37 °C (20 min) → 80 °C (20 min)<br>Ligation: 25 °C (30 min) → 65 °C (10 min) |
| BioBricks | Digestion: 37 °C (20 min) → 80 °C (20 min)<br>Dephosphorylation of backbone DNA: 37 °C (30 min) → 80 °C (5 min)<br>Ligation: 25 °C (30 min) → 65 °C (10 min) |
| Bri2Mo First piece | Digestion: 37 °C (20 min) → 80 °C (20 min)<br>Dephosphorylation of backbone DNA: 37 °C (30 min) → 80 °C (5 min)<br>Ligation: 25 °C (30 min) → 65 °C (10 min) |
| Bri2Mo Additional piece | Digestion: 37 °C (20 min) → 80 °C (20 min)<br>Dephosphorylation of backbone DNA: 37 °C (30 min) → 80 °C (5 min)<br>Ligation: 25 °C (30 min) → 65 °C (10 min) |
| Bri2Mo mutagenic PCR | Initial DNA denaturalization: 98°C (30 sec)<br>31 cycles:<br>DNA denaturalization: 98 °C (10 sec)<br>Primer annealing: 69 °C (20 sec)<br>Extension: 72 °C (37 sec)<br>Final extension: 72 °C (2 min) |

#### Mo2Bri operation

**Table S5. One-pot cloning.** Detailed Mo2Bri reactants for DNA digestion and dephosphorylation (step 1 in Figure 2A). Note that Mo2Bri one-pot reactions also include ligation (step 2 in Figure 2A). Note also that rSAP is added right after inactivating the digestion of backbone DNA.

| Reactants | One-pot (37°C & 42°C) | Sequential (rSAP- & rSAP+) |
| --- | --- | --- |
| Molecular-grade water | Up to 10 µL | Up to 10 µL |
| Backbone DNA | 50 fmol | 100 ng |
| Bri2Mo Insert DNA | 50 fmol | 100 ng |
| BsmBI-v2 | 5 units | 1.1 units |
| 10x r3.1 buffer | - | 1 µL |
| T4-DNA ligase | 200 units | - |
| 10x T4-DNA ligase buffer | 1 µL | - |
| rSAP | - | 1 unit (only for rSAP+) |

**Table S6. Two-pot Cloning.** Detailed sequential Mo2Bri reactants for DNA ligation (step 2 in Figure 2A).

| Reactants | Sequential (rSAP+ and rSAP -) |
| --- | --- |
| Molecular-grade water | 2.25 µ |
| Digested backbone DNA | 0.5 µL |
| Digested Bri2Mo Insert DNA | 1.5 µL |
| T4-DNA ligase | 100 units |
| 10x T4-DNA ligase buffer | 0.5 µL |

#### Bri2Mo operation

**Table S7.** Detailed Bri2Mo\_1 reactants for DNA digestion (step 1 in Figure 2A).

| Reactants | Bri2Mo_2 |
| --- | --- |
| Molecular-grade water | Up to 10 μL |
| DNA acceptor (Backbone with MCS2 plasmid (ii) in Fig 2A) | 100 ng |
| Backbone DNA donor (Backbone with insert, plasmid (i) in Fig 2A) | 100 ng |
| BsmBI-v2 | 1.1 units |
| 10x r3.1 buffer | 1 μL |
| rSAP | 1 unit |

**Table S8.** Detailed Bri2Mo\_2 reactants for DNA ligation (step 2 in Figure 2A).

| Reactants | Bri2Mo_2 |
| --- | --- |
| Molecular-grade water | 2.25 μL |
| BsmBI-digested DNA acceptor | 0.5 μL |
| BsmBI-digested DNA donor | 1.5 μL |
| T4-DNA ligase | 100 units |
| 10x T4-DNA ligase buffer | 0.5 μL |

**Table S9.** Detailed BioBrick digestion and dephosphorylation reactants (step 1 in Figure 3A and 3 in Figure 3C).

| Reactants | Bri2Mo_2 |
| --- | --- |
| Molecular-grade water | Up to 10 μL |
| Backbone DNA | 100 ng |
| BioBrick insert DNA | 100 ng |
| EcoRI-HF | 2.2 units |
| SpeI-HF | 2.2 units |
| XbaI | 2.2 units |
| PstI-HF | 2.2 units |
| 10x rCutSmart buffer | 1 μL |
| rSAP | 1 unit |

**Table S10.** Detailed BioBrick ligation reactants (step 2 in Figure 2A and step 3 in Figure 3C).

| Reactants | Bri2Mo_2 |
| --- | --- |
| Molecular-grade water | 2.25 μL |
| Digested backbone DNA | 0.5 μL |
| Digested BioBrick insert DNA | 1.5 μL |
| T4-DNA ligase | 100 units |
| 10x T4-DNA ligase buffer | 0.5 μL |

**Table S11.** Detailed Bri2Mo\_3 (erasing) site mutagenic PCR reactants.

| Reactants | Bri2Mo_3 mutagenic PCR |
| --- | --- |
| Molecular-grade water | 31.5 μL |
| Template DNA (1 pg - 10 ng) | 1 μL |
| 10 mM dNTPs | 1 μL |
| 10 μM Erasing Forward primer | 2.5 μL |
| 10 μM Erasing Reverse primer | 2.5 μL |
| Q5-DNA polymerase | 1 unit |
| 5x Q5-DNA polymerase buffer | 10 μL |

#### **SPECTRA code depiction.**

Briefly, images were subjected to automated region-of-interest (ROI) detection via Hough Circle Transform, followed by a 15-pixel morphological erosion mask to eliminate plastic edge reflections using OpenCV (2). For colony segmentation, a Top-hat morphological filter (31 × 31 elliptical kernel) was applied to the grayscale image for background subtraction, followed by Euclidean distance transformation and Watershed segmentation to accurately resolve adjacent colonies, implemented via the scikit-image library (3). These traditional computer vision algorithms remain highly competitive and are widely integrated into modern microbiological pipelines (4). Phenotypic classification (e.g., GFP+, RFP+, or non-fluorescent) was executed using a supervised machine learning approach. A linear Support Vector Machine (SVM) classifier (5), implemented via scikit-learn (6), was trained in situ using few-shot user annotations. To ensure robustness against lighting variations, the classifier evaluated a 6-dimensional spectral feature space extracted from the BGR color matrices (R, G, B, R/G, G/B, R/B). Only detections with an SVM probability score >0.65 and passing a dynamic size-exclusion filter (40–250% of the median valid colony area) were initially quantified. Following the automated analysis, a human-in-the-loop curation step was incorporated. This enabled users to visually inspect the AI-generated segmentations, manually correct potential misclassifications. This code is available for the community at GitHub (https://github.com/alonso1310/SPECTRA).

#### **In-house protocols**

##### Preparation of chemically competent cells

A fresh preculture was prepared by inoculating the NEB 10-beta *E. coli* strain into 4 mL of LB medium with streptomycin. The culture was incubated overnight at 37°C with shaking at 280 rpm.

The following day, 1 mL of the overnight preculture was inoculated into two separate flasks, each containing 100 mL of sterile LB medium without antibiotic. The cultures were incubated at 37°C with shaking at 280 rpm until they reached an OD600 of 0.4–0.6.

Once the desired optical density was reached, both cultures were equally divided into sterile 50 mL centrifuge tubes and centrifuged at 5,000 × g for 10 minutes at 4°C. After centrifugation, the supernatant was carefully discarded, and each bacterial pellet was gently resuspended in 15 mL of ice-cold 0.1 M CaCl<sub>2</sub> solution. The cell suspensions were then incubated on ice for 10 minutes.

Following this incubation, the cells were centrifuged again at 5,000 × g for 5 minutes at 4°C, and the supernatant was carefully discarded. The pellets from two of the tubes were gently resuspended in 3 mL of ice-cold 0.1 M CaCl<sub>2</sub> containing 15% (v/v) sterile glycerol, avoiding excessive pipetting to minimize cell damage. Each cell suspension was then transferred to one of the remaining tubes containing a bacterial pellet, which was subsequently resuspended, thereby doubling the cell concentration in the final suspension.

The resulting competent cell suspension was aliquoted into 150 µL portions in pre-chilled sterile 1.5 mL tubes, which were kept on dry ice throughout the process. Finally, the aliquots were immediately stored at –80°C in a properly labeled storage box until use.

#### Preparation of Solutions

For the 0.1 M $\text{CaCl}_2$ solution, sterile molecular grade water was used, reaching a volume of 50 mL. The 0.1 M $\text{CaCl}_2$ solution supplemented with 15% (v/v) glycerol was prepared fresh on the day of use and maintained at 4°C before and throughout the procedure. To prepare 30 mL of this solution, 4.5 mL of sterile 100% glycerol was mixed with 25.5 mL of 0.1 M $\text{CaCl}_2$. The solution was kept on ice until required.

#### Protocol of bacterial transformation

Chemically competent *E. coli* NEB 10-beta cells were thawed on ice and maintained at 0-4°C throughout the transformation procedure. Plasmid DNA was also kept on ice to ensure that both the DNA and competent cells remained at the same temperature prior to transformation.

For each transformation, 50 $\mu\text{L}$ of competent cells were transferred into a sterile 1.5 mL microcentrifuge tube, and 2.5 $\mu\text{L}$ of cloning DNA were added directly to the cell suspension. The mixture was gently resuspended by pipetting. The DNA–cell mixture was then incubated on ice for 20 minutes.

Following the incubation on ice, the cells were subjected to heat shock at 42°C for 45 seconds using a preheated thermomixer or heating block. Immediately after heat shock, the tubes were returned to ice and incubated for 5 minutes.

After recovery on ice, 200 $\mu\text{L}$ of pre-warmed 2YT medium were added to each transformation mixture, resulting in a final volume of 250 $\mu\text{L}$. The cells were incubated at 37°C with shaking at 180 rpm for 1 hour.

Afterwards, between 50 and 100 $\mu\text{L}$ of the cells were spread onto LB agar plates supplemented with the appropriate antibiotic using a sterile spreader or sterile glass beads. The plates were allowed to absorb the inoculum for approximately 5 minutes before being incubated inverted at 37°C overnight. Transformed colonies were subsequently selected based on their growth on the appropriate antibiotic-containing medium and their expected phenotypic characteristics.
